# Emerging experience-dependent dynamics in primary somatosensory cortex reflect behavioral adaptation

**DOI:** 10.1101/2021.01.29.428886

**Authors:** Christian Waiblinger, Megan E McDonnell, Peter Y Borden, Garrett B Stanley

**Author notes:** Correspondence: Garrett B Stanley, Coulter Department of Biomedical Engineering, Georgia Institute of Technology & Emory University, 313 Ferst Drive, Atlanta GA 30332-0535, USA.

## Abstract

Behavioral experience and flexibility are crucial for survival in a constantly changing environment. Despite evolutionary pressures to develop adaptive behavioral strategies in a dynamically changing sensory landscape, the underlying neural correlates have not been well explored. Here, we use genetically encoded voltage imaging to measure signals in primary somatosensory cortex (S1) during sensory learning and behavioral adaptation in the mouse. In response to changing stimulus statistics, mice adopt a strategy that modifies their detection behavior in a context dependent manner as to maintain reward expectation. Surprisingly, neuronal activity in S1 shifts from simply representing stimulus properties to transducing signals necessary for adaptive behavior in an experience dependent manner. Our results suggest that neuronal signals in S1 are part of an adaptive framework that facilitates flexible behavior as individuals gain experience, which could be part of a general scheme that dynamically distributes the neural correlates of behavior during learning.

## INTRODUCTION

Survival in a dynamically changing sensory environment requires a high degree of behavioral flexibility and experience. While much is known about the origin and processing of sensory signals in mammalian brains^1–3^, far less is understood about how resultant behavioral strategies are formed with practice and experience, and the role of primary sensory areas in this output process. Studies have investigated the role of the primary sensory cortex in visual^4–6^, auditory^7–10^ and somatosensory behaviors^11, 12^. There are varying views on whether primary sensory cortex simply relays an ascending sensory signal or plays a deeper role in decision making and behavior.

It is likely that primary cortical signals are highly dynamic and context driven, depending on the implemented behavioral paradigm. Signals in primary somatosensory cortex can: enhance stimulus selectivity with behavioral training^13^, fluctuate according to the behavioral state^14^, or even remap depending on downstream signals^15^. Neuronal signals in primary sensory areas may be highly dynamic, context or experience dependent, and part of an adaptive framework.

Here, we investigate perceptual capabilities of the primary somatosensory cortex (S1) during learning and behavioral adaptation, using the highly conserved mouse vibrissa system. We hypothesize that signals in S1 not only represent the strength of a sensory input but also play a key role in the transformation of context dependent behaviors. To test this, we designed a series of psychophysical experiments evaluating behavioral performance and neuronal activity during: 1) gradual learning of a basic detection task and 2) adaptation to repetitive changes in sensory contingencies. To repeatedly measure signals of large neuronal pools across both training stages, we performed chronic wide-field imaging of S1 activity with the genetically encoded voltage indicator (GEVI) “ArcLight” ^16, 17^ in behaving mice.

During learning of the basic task, S1 sensitivity is mostly stimulus driven and uncorrelated with gradual changes in behavioral performance. Interestingly, basic detection was not abrogated by S1 inactivation through lesioning. In contrast, our results further reveal that S1 activity correlates with behavioral adaptation, following long-term exposure to changing sensory stimulus statistics. Mice adopt a strategy that modifies their behavior in a way as to maintain reward in the face of these changes: once an animal is trained to adapt to a change in stimulus statistics, neuronal activity dynamically shifts between changes in S1 sensitivity and decision criterion downstream. S1 inactivation through lesioning disrupted the adaptive behavior, suggesting the S1 primary cortex is necessary for an adaptive response to dynamic stimuli.

Our findings suggest a translation of these context dependent changes between different brain structures along the hierarchy, where S1 is not only producing primary neuronal signals in response to tactile input, but also transducing signals necessary for adaptive behavior strategies in a dynamically changing environment.

## MATERIALS AND METHODS

### Animals, surgery, and general procedures for behavioral testing

All experimental and surgical procedures were approved by the Georgia Institute of Technology Institutional Animal Care and Use Committee and were in agreement with guidelines established by the NIH. Subjects were 13 male mice (C57BL/6, Jackson Laboratories), aged 4-6 weeks at time of implantation. The basic procedures of virus delivery, head-plate preparation and cortical imaging exactly followed the ones published in a recent paper^16^. In the following text, only procedures pertaining to the specific procedures established here are described in detail.

### Virus delivery

At least 4 weeks prior to experimentation, mice were anesthetized using isoflurane, 3-5 % in a small induction chamber, and then placed on a heated platform (FHC, Inc.) to maintain body temperature with a stereotaxic nose cone to maintain anesthesia. During the surgery, the anesthesia levels were adjusted to 1-1.5 % to achieve ∼1/s breathing rate in mice. For virus delivery, 3 small craniotomies (burr holes of 0.7 mm diameter) were created over the barrel field of S1 according to stereotaxic measurements taken from the bregma ([1 x 3 mm, 3 x 3 mm, 3 x 1 mm] bregma x lateral). The virus was loaded into a neural syringe (Hamilton Neuros Syringe 700/1700). The injection needle was initially lowered to 1000 µm below the pia surface for pre-penetration and then retracted to the target depth of 500 μm, using a 10 μm resolution stereotaxic arm (Kopf, Ltd.). Following a 1-minute delay to allow for tissue relaxation, each animal was injected with 1.5 µl of adeno-associated virus (AAV)1-hsyn1-ArcLight-D-WPRESV40 (UPenn Viral Vector Core, AV-1-36857P) at a flow rate of 0.05 μL⁄min (0.5 μl each, for three injections). After injection, the needle remained in place for an additional 5 minutes before slowly being removed from the brain. The craniotomies were left to close naturally. The skull was sealed by suturing the skin. Throughout the experiment, sterile techniques were used to keep the injection area clean and free from infection. Additionally, opioid and non-steroidal anti-inflammatory analgesic were administered (SR-Buprenorphine 0.8-1 mg/kg, SC, pre-operatively and Ketoprofen 5-10 mg/kg, IP, post-operatively).

### Head plate Implantation

After at least 4 weeks post injection, a metal head-plate was secured to the skull in order to reduce vibration and allow head-fixation during imaging and behavior experiments. Following anesthetization and analgesia, a large incision was made over the skull. The connective tissue and muscles surrounding the skull were removed using a fine scalpel blade (Henry Schein #10). The custom titanium head-plate formed an open ring (half-moon shape with an inner radius of 5 mm) and was placed on top of the two hemispheres with an extended bar above the cerebellum (∼10 mm, perpendicular to the midline of the skull). The extended bar was designed to attach to a stainless steel holder, serving the purpose of stable head-fixation. The head-plate was attached to the bone using a three stage dental acrylic, Metabond (Parkell, Inc.). The Metabond was chilled using ice, slowly applied to the surface of the skull, and allowed to cure for 5-10 minutes. After securing the head-plate, the skull was cleaned and covered with a thin layer of transparent Metabond. During preparation for histological validation, the head-plate could not be separated from the attached skull and the brain was extracted by removing the lower jaw. The final head-plate and dental acrylic structure additionally created a well for mineral oil that helped maintain skull transparency for the upcoming imaging sessions. Mice were allowed to recover for at least 7 days before habituation training. After wound healing, subjects were housed together with a maximum number of three in one group cage and kept under a 12/12 h inverted light/dark cycle.

### Cortical Lesions

After the collection of at least 5 sessions of psychometric data and the mapping of spatial activation using GEVI imaging, lesions of the barrel field in S1 were performed as a control experiment (n = 4 mice). Animals were given water *ad libitum* for the day preceding the procedure. Following anesthetization, a small craniotomy (burr hole of 0.7 mm diameter) was created over the area in S1 with the highest extent of activation from the GEVI map. The neurotoxin ibotenic acid^18^ (aablocks, AA003BBF) at a concentration of 10 mg/mL was loaded into a neural syringe (Hamilton Neuros Syringe 700/1700). The injection needle was lowered to 1000 μm depth for pre-penetration, then retracted to a depth of 500 μm, using a 10 μm resolution stereotaxic arm (Kopf, Ltd.). Following a delay of 5 minutes for tissue relaxation, animals were injected with 0.25 μL of ibotenic acid at a flow rate of 0.05 μL/min. After injection, the needle remained in place for 5 additional minutes before being slowly retracted from the brain. The craniotomy was sealed using the dental acrylic Metabond (Parkell, Inc.), and the silicone elastomer Kwik-Cast (Kwik-Cast Sealant World Precision Instruments), for protection. Animals were allowed to recover for 1-3 days until activity and body weight were normal, and then behavioral testing and GEVI imaging continued.

### Whisker Stimulation

Precise whisker deflections were performed using a calibrated galvo-motor (galvanometer optical scanner model 6210H, Cambridge Technology) as described in a previous study^19^. The opening of the rotating arm was narrowed with dental cement to prevent whisker motion at the point of insertion. The rotating arm contacted a single whisker on the right of the mouse’s face at 5 mm (±1 mm tolerance) distance from the skin, and thus, directly engaged the proximal whisker shaft, largely overriding bioelastic whisker properties. Distance and angle between stimulator and whisker were systematically measured with a camera before each session to ensure that stimulation was consistent. All the remaining whiskers were trimmed to prevent them from being touched by the rotating arm. Across mice, different whiskers were chosen (C1, D1, D2 or E2), but the same whisker was used throughout sessions within each mouse Voltage commands for the actuator were programmed in Matlab and Simulink (Ver. 2015b; The MathWorks, Natick, Massachusetts, USA). A stimulus consisted in a single event, a sinusoidal pulse (half period of a 100 Hz sine wave, starting at one minimum and ending at the next maximum). The pulse amplitudes used (A = [0, 1, 2, 4, 8, 16] °, correspond to maximal velocities: V _max_ = [0 314 628 1256 2512 5023] °/s) or mean velocities: V _mean_ = [0 204 408 816 1631 3262] °/s) and were well within the range reported for frictional slips observed in natural whisker movement^20, 21^.

### Cortical GEVI Imaging

ArcLight transfected mice were chronically imaged through the intact skull using a wide-field fluorescence imaging system to measure cortical spatial activity (MiCAM05-N256 Scimedia, Ltd.). Figure 1a shows the experimental apparatus and schematically describes the wide-field fluorescence microscope. During all imaging experiments, mice were awake and head-fixed. The head-plate was used as a well for mineral oil in order to keep the bone surface wet and maintain skull transparency. The skull was covered with a silicone elastomer (Kwik-Cast Sealant World Precision Instruments) between imaging sessions for protection. The barrel cortex was imaged using a 256 x 256 pixel CMOS camera (Scimedia Model N256CM) at 200 Hz with a pixel size of 69 μm x 69 μm and an active imaging area of 11.1 x 11.1 mm, given a magnification of 0.63. Note, this resolution does not consider the scattering of the light in the tissue. During experimental imaging, the illumination excitation light was left continuously on. The entire cortical area was illuminated at 465 nm with a 400 mW/cm^2^ LED system (Scimedia, Ltd.) to excite the ArcLight fluorophore. The excitation light was further filtered (cutoff: 472-30 nm bandpass filter, Semrock, Inc.) and projected onto the cortical surface using a dichroic mirror (cutoff: 495 nm, Semrock, Inc.). Collected light was filtered with a bandpass emission filter between wavelengths of 520-35 nm (Semrock, Inc.). The imaging system was focused at ∼300 μm below the cortical surface to target cortical layer 2/3. The first imaging session was used for identifying the barrel field in the awake and naïve animal at least four weeks after ArcLight viral injection. The barrel field was mapped by imaging the rapid response to a sensory stimulus given to a single whisker (A = 4-16°, or mean velocities respectively: V = 816-3262°/s). We used two criteria to localize and isolate the barrel field: standard stereotaxic localization (∼3 mm lateral, 0.5-1.5 mm caudal from bregma) and relative evoked spatial and temporal response (visible evoked activity 20-25 ms after stimulation). A single whisker was chosen if it elicited a clear response within the barrel field. All subsequent imaging experiments were centered on the same exact location and the same whisker was chosen in repeated sessions for a given animal. Figure 1b shows the characteristic spread of ArcLight expression in an example coronal brain section. The same hemisphere is shown *in vivo* in Figure 1c with spontaneous fluorescence activity in S1 at the beginning of a behavioral session. Spontaneous and sensory evoked activity in S1 was imaged every trial, 2 s before and 2 s after the punctate whisker. Figure 1d shows frames with typical fluorescence activity patterns recorded at a framerate of 200 Hz and shown separately for three different stimulus amplitudes (rows). The calculation of the ΔF/F_0_ metric, region of interest (ROI) and data processing is described under “Imaging analysis”.

**Figure 1.**
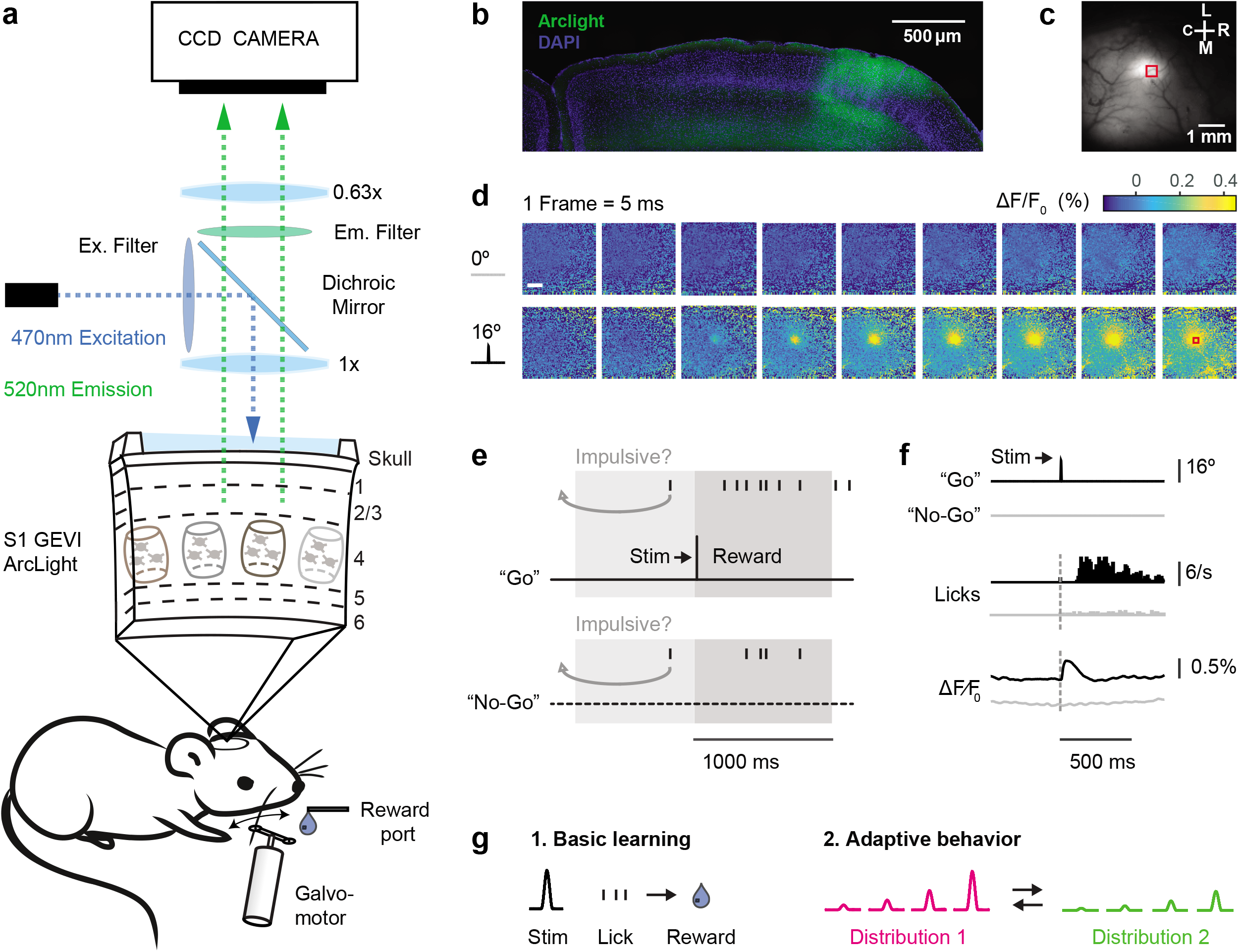
Imaging and behavior methods. **a,** Top: Schematic of the imaging system. The GEVI “ArcLight” is expressed in superficial layers of S1. Bottom: Schematic of the behavior setup. **b,** Confocal image of a coronal mouse brain section (100 μm, right hemisphere) showing the characteristic spread of ArcLight in S1. Blue: DAPI staining, Green: Arclight fluorescence. **c,** Top view of the same hemisphere in vivo showing ArcLight fluorescence in S1. **d,** Images of the same hemisphere in a trained mouse with ArcLight fluorescence color coded. Punctate stimuli (16 degree whisker angle) or catch trials (0 degree) were presented. Frames were captured at 200 Hz and depicted from stimulus onset onward. Each frame is normalized to the frame at stimulus delivery (ΔF⁄F_0_ = F-F_0_/F_0_). Shown are fluorescence responses averaged over an example imaging session (n=24 trials/condition). A region of interest (red box, 434 x 434 µm) is centered at the peak fluorescence to extract and average voltage traces for further analysis. Scale bar: 1 mm. **e,** Go/No-Go detection task. A punctate stimulus (10 ms) has to be detected by the mouse with an indicator lick to receive reward. Reward is only delivered in hit trials. Impulsive licks in a 2 s period before trial onset are mildly punished by a time-out triggering a new inter-trial-interval (4-10 s, gray arrow). **f,** Example traces of Go and No-Go trials (n=284 each). Top: Readouts from the galvo sensor (16 degree stimulation or 0 degree catch trial). Middle: Lick response histograms from a trained mouse. Bottom: Average voltage response from the region of interest of S1. **g,** Different task versions under investigation. 1. Basic learning: Learning of the Go/No-Go task. 2. Adaptive behavior: Once animals have learned the basic task, they are challenged with multiple stimulus amplitudes and changes in the statistics of the stimulus distribution.

### Behavioral paradigm and training

During successive days of behavioral testing, water intake was restricted to the experimental sessions where animals were given the opportunity to earn water to satiety. Testing was paused and water was available *ad libitum* during 2 days a week. Body weight was monitored daily, and was typically observed to increase or remain constant during training. In some cases, the body weight dropped slightly across successive training days due to a higher task difficulty. If the weight dropped for more than ∼5g, supplementary water was delivered outside training sessions to maintain the animal’s weight. 1-2 training sessions were usually conducted per day comprising 50-250 trials. During behavioral testing a constant auditory white background noise (70 dB) was produced by an arbitrary waveform generator to mask any sound emission of the galvo-motor-based whisker actuator. All mice were trained on a standard Go/No-Go detection task (Fig. 1e) employing a similar protocol as described before ^22–26^. In this task, the whisker is deflected at intervals of 4-10 s (flat probability distribution) with a single pulse (detection target). A trial was categorized as a “hit” if the animal generated the “Go” indicator response, a lick at a water spout within 1000 ms of target onset. If no lick was emitted the trial counted as a “miss”. In addition, catch trials were included, in which no deflection of the whisker occurred (A = 0°) and a trial was categorized as a “correct rejection” if licking was withheld (“No-Go”). However, a trial was categorized as a “false alarm” if random licks occurred within 1000 ms of catch onset. Premature licking in a 2 s period before the stimulus was mildly punished by resetting time (“time-out”) and starting a new inter-trial interval of 4-10 s duration, drawn at random from a flat probability distribution. Note these trial types were excluded from the main data analysis.

The first step of behavioral training was systematic habituation to head-fixation and experimental chamber lasting for about one week. During the second training phase, a single whisker deflection with fixed amplitude was presented interspersed by catch trials (P_stim_ = 0.8, P_catch_ = 0.2). Immediately following stimulus offset, a droplet of water became available at the waterspout to condition the animal’s lick response thereby shaping the stimulus-reward association. Once subjects showed stable and immediate consumption behavior (usually within 1-2 sessions), water was only delivered after an indicator lick of the spout within 1000 ms, turning the task into an operant conditioning paradigm in which the response is only reinforced by reward if it is correctly emitted after the stimulus. Subsequent experiments were performed systematically and the behavioral performance was measured with simultaneous GEVI imaging. The different experiments are described in detail in the following section.

### Basic learning

Learning was studied once an animal had entered the operant phase of training after the basic habituation procedure. From this point forward, experiments were conducted with equal conditions across sessions and without manual interference by the experimenter. To assess differences in learning based on stimulus strength, animals were separated into two groups: group 1 (n = 3) receiving only one low amplitude stimulus and catch trials (A = [4 0] °, Fig. 2) and group 2 (n = 3) receiving only one high amplitude stimulus and catch trials (A = [16 0] °, Suppl. Fig. 2). Performance metrics are described under *data analysis and statistics*.

**Figure 2.**
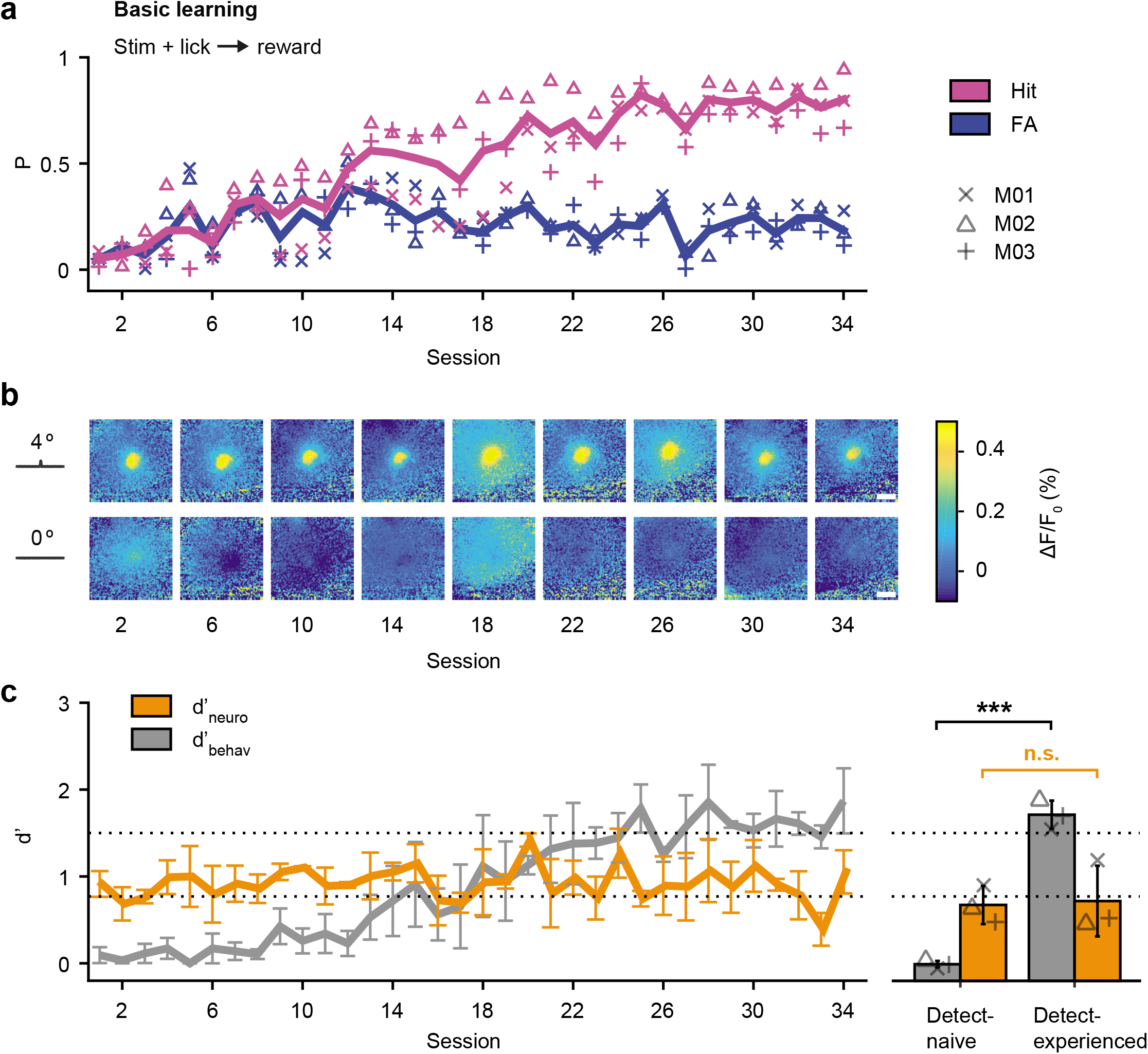
Basic Learning and sensory signals in S1. **a,** Learning curve for 3 mice trained on the basic Go/No-Go detection task with a weak stimulus (4 degree). Performance is expressed as the probability of a hit (lick after stimulus) and false alarm (lick after no stimulus) for a given session. Individual mice are represented by symbols, and the mean by bold lines. **b,** Fluorescent activity in S1 from an example mouse during learning of the task. Shown are frames at the peak response to the 4 degree stimulus, catch trials are shown below. Scale bar: 1 mm. **c,** Dprime metrics for both behavioral and neuronal data during learning of the task. The d’_behav_ was calculated from the observed hit rate and false alarm rate for a given training session. The d’_neuro_ was calculated by comparing single trial distributions of all evoked signal peaks (maximum % ΔF⁄F_0_ within 100 ms post stimulus) with the corresponding noise distributions when no stimulus was present. Shown are means across mice (n=3) for a given session, errorbars represent bootstrapped estimates of 95% confidence limits (nBoot=1000 repetitions). The dotted lines separate performance into “detect-naive” (d’<0.8), and “detect-experienced” (d’>1.5). The right panel shows the same data separated for individuals (symbols) before and after learning. Bars represent means across mice, errorbars represent SD (n=3). * P < 0.05; ** P < 0.01; *** P < 0.001; ‘n.s.’ not significant, two-sided Wilcoxon rank-sum test.

### Adaptive behavior

After mice had learned the basic detection task, the psychometric curve was measured in a subgroup of animals (n = 10) using the method of constant stimuli, which entails the presentation of repeated stimulus blocks containing multiple stimulus amplitudes. On a single trial, one out of multiple possible stimulus amplitudes was presented after a variable time interval (4-10 s), each with equal probability (uniform distribution, p = 0.2). A stimulus block consisted of a trial sequence comprising all stimuli and a catch trial in pseudorandom order (e.g., each type once per block). A behavioral session consisted of repeated stimulus blocks until the animal disengaged from the task, i.e. when it did not generate lick responses for at least an entire stimulus block. Therefore, the chosen stimulus occurred repetitively but randomly within a session. A condition was always kept constant within and across multiple behavioral sessions before the task was changed. A “switch” was defined as the transition between two sessions with different conditions. In the “high range” condition, four stimulus amplitudes plus catch trial were used (A = [0, 2, 4, 8, 16] °) and presented in multiple successive sessions. Following this, four new stimulus amplitudes were presented (A = [0, 1, 2, 4, 8] °) forming the “low range” condition. Both stimulus distributions shared two of the three stimulus amplitudes; however, the largest stimulus amplitude of the high range condition (A = 16 °) was not part of the low range, and vice versa, the smallest amplitude of the low range condition (A = 1 °) was not part of the high range. Conditions were always changed in alternating fashion to test the reversibility of adaptation (e.g. high-low-high or vice versa) and multiple switches were performed to test adaptation at different training stages. The different timescales are summarized as follows:

### Number of trials

the number of trials were variable since each session was stopped after an animal missed an entire set of stimuli (i.e. A = [0, 2, 4, 8, 16] ° for the high range). The reason for this implementation is to ensure that the data are not affected by the animal’s potential satiation on any given day. As shown in Figure 3c, the number of trials per session averages out to approximately one hundred trials (n = 101 for high range and n = 104 for low range).

**Figure 3.**
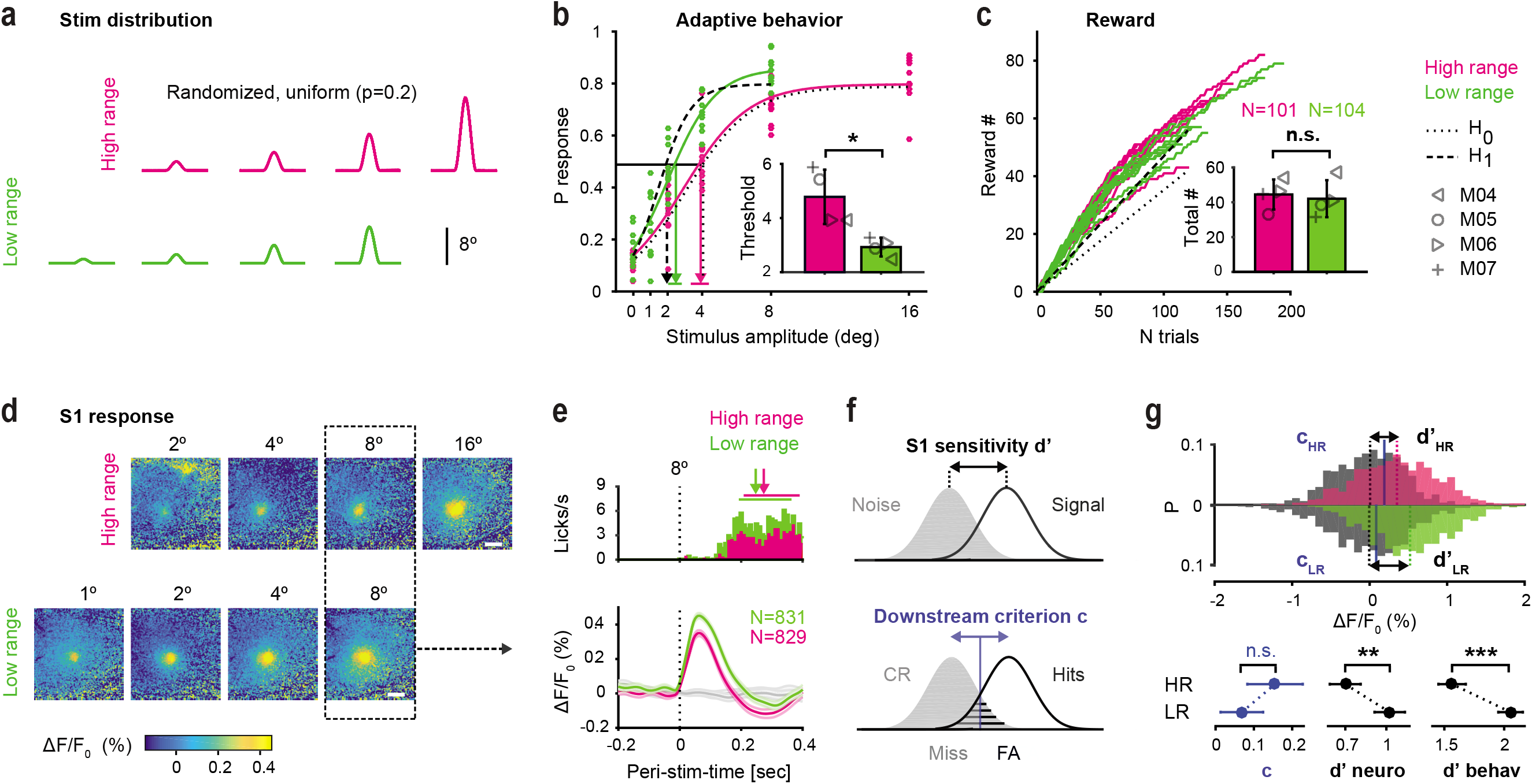
Adaptive behavior and S1 responses. **a,** Manipulation of stimulus distribution range. Every stimulus and catch trial (not shown) is presented with equal probability (p=0.2). The design involves amplitudes common to both high range (magenta) and low range (green) conditions. **b,** Psychometric curves and response thresholds for an example animal working on both conditions. Each dot corresponds to response probabilities from a single session. Solid curves are logistic fits to the average data (n=10-11 sessions). Dotted line is a hypothetical curve assuming no change in performance (H_0_). Dashed line is a hypothetical curve assuming a change in performance to maintain reward (H_1_) when switching from high range to low range stimulus distributions. Response thresholds are shown as vertical lines with 95% confidence limits. Inset. Response thresholds of all mice (grey symbols). Bars represent means across mice with SD (n=4). **c,** Number of rewards (correct trials) accumulated by the same animal from b. Each line corresponds to one session. Inset. Average total reward number per session for each mouse. The average number of trials is shown on top. Figure conventions are the same as in b. **d,** Frames of evoked cortical fluorescence activity from two example sessions, one with a high range stimulus distribution (top) and the other with a low range stimulus distribution (bottom). The frames are aligned for amplitudes common to both datasets. Data with a deflection angle of 8 degree was chosen for further analysis (outlined box). **e,**Temporal fluorescent signal in response to 8 degree stimulation, extracted from the region of interest and averaged across sessions and mice. PSTHs of behavioral lick responses are shown on top. Median reaction times (first lick post stimulus) are shown as arrows, 25-75 percentiles as horizontal lines (n=829-831 trials). **f,** Observer model based on signal detection theory. Behavioral adaptation can either be induced by changes in sensitivity (d’) intrinsic to S1, changes in decision criterion by a downstream observer, or both. **g,** Top: Distributions of evoked trials (signal) and catch trials (noise) computed separately for the high (HR) and low range (LR) condition. Dashed lines indicate mean ΔF/F_0_, black arrows indicate dprime values. Blue lines indicate criterion values as computed from ROC curves. Bottom: Criterion and dprime metrics (neuronal and behavioral). Shown are bootstrapped estimates of means and 95% confidence limits (n=38 (LR) or n=41 (HR) sessions, n=4 mice, nBoot=1000 repetitions). * P < 0.05; ** P < 0.01; *** P < 0.001; ‘n.s.’ not significant, two-sided Wilcoxon rank-sum test or Kruskal-Wallis test.

### Number of sessions

Within a block there are 5-11 daily sessions depending on the animal. However, each animal experienced a matched number of sessions across blocks (e.g. n = 5 for high range and n = 5 for low range).

### Number of blocks

Depending on the animal, 3-6 blocks were performed with alternating conditions either starting with the high range or the low range condition (resulting in 2-5 switches).

### Histology

Upon completion of experiments, transcardiac perfusion was performed on all animals with a 4 % paraformaldehyde solution in phosphate-buffered saline. The head plates were carefully removed from the skulls, and the brains were extracted and post-fixed for 2-24 hours depending on the success of the perfusion. Brains were then sectioned at 100µm thickness on a vibratome (Leica VT1000S). The brain sections were stained with Cytochrome C oxidase and DAPI for histological analysis of lesion extent or ArcLight expression. Sections were mounted and imaged using a wide field color microscope (Zeiss Axio Observer Z1).

### Data analysis and statistics

*Behavior*. The learning curve was measured by calculating a dprime, *d*′_*behav*_, from the observed hit rate and false alarm rate of each training session

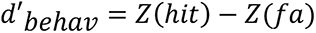

where the function Z(p), p ∈ [0,1], is the inverse of the cumulative distribution function of the Gaussian distribution. A criterion of d’ = 2.3 (calculated with *p*(*hit*) = 0.95 and *p*(*fa*) = 0.25) was used to determine the end of the basic learning period and learning progress was separated into three equally distributed stages, “detect-naive” (*d*’ = 0-0.8), “detect-interim” (*d*’ = 0.8-1.5) and “detect-experienced” level (*d*’ = 1.5-2.3). Psychometric data were assessed as response-probabilities averaged across sessions within a given stimulus condition.

Psychometric curves were fit using Psignifit^27–29^. Briefly, a constrained maximum likelihood method was used to fit a modified logistic function with 4 parameters: α (the displacement of the curve), β (related to the inverse of slope of the curve), γ (the lower asymptote or guess rate), and λ (the higher asymptote or lapse rate) as follows:

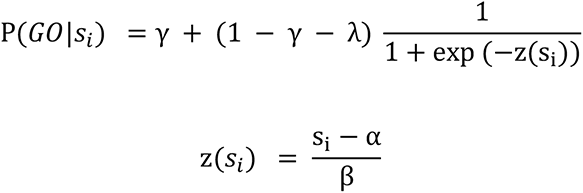

where *s_i_* is the stimulus on the i^th^ trial. Response thresholds were calculated from the average psychometric function for a given experimental condition using Psignifit. The term “response threshold” refers to the inverse of the psychometric function at some particular performance level with respect to the stimulus dimension. Throughout this study, we use a performance level of 50% (probability of detection of 0.5). Statistical differences between psychophysical curves were assessed using bootstrapped estimates of 95% confidence limits for the response thresholds provided by the Psignifit toolbox. To assess the effects of the lesion on behavior, the psychometric curves and response thresholds were compared in lesion animals, from before and after the lesion, and between high- and low-range stimuli. Kruskal-Wallis or rank sum tests were used to compare significance of the lesion effects on response threshold in lesioned versus healthy animals.

### Reward accumulation

Let the stimulus amplitude delivered on the i^th^ trial be denoted as *s_i_*, the corresponding reward as *r_i_*, (since *r_i_* is a fixed value, it can just be termed r) and the accumulated reward for N trials as *R_N_*. Over N trials, the expected accumulated reward is

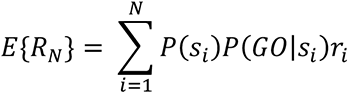

where *P*(*s_i_*) comes from the experimentally controlled stimulus distribution, *P*(*GO*|*s_i_*) is the probability of a positive response (or “Go”) for the given stimulus amplitude, and *E*{ } denotes statistical expectation.

We considered the null hypothesis of this behavioral paradigm to be that animals do not adapt their behavior in response to an experimentally forced change in stimulus distribution and thus operate from the same psychometric curve (represented as dotted curves in Fig. 3b). Note, this corresponds to the same curve but for a different range of stimuli (*P*(*GO*|*s_i_*) across different *s_i_*. For example, in moving from the high range to the low range stimulus condition, this would result in a decrease in the total accumulated reward for the same number of trials.

As an alternative hypothesis, one possible strategy the animal could take in response to a change in the stimulus distribution would be to adjust behavior to maintain the same amount of accumulated reward during a session. For example, in moving from high range to low range stimuli, the accumulated reward would be assumed fixed, and we can determine a new set of probabilities *P*(*GO*|*s_i_*) that define an adapted psychometric function. Note that there is not a unique solution, but one simple possibility is that the original psychometric function maintains the same asymptotes (γ and λ) and false alarm rate but is compressed, with a decrease in response threshold and an increase in slope to maintain the same total accumulated reward. We denote this situation as our hypothetical psychometric function, represented as dashed curves in Figure 3b.

### Imaging analysis

All voltage imaging data was analyzed using custom written image-analysis software (MATLAB 2015a, Mathworks, Inc.). The specific methods of processing the ArcLight raw fluorescence signal and basic data analysis followed those of a recent study from our laboratory^16^. Briefly, raw images were loaded and converted from the proprietary file format of the imaging system using custom scripts. Due to the natural decay of the fluorescent signal caused by photo bleaching, each trial was first normalized to a baseline and reported as a percent change in fluorescent activity (% ΔF/F_0_). The ΔF/F_0_ measurement was calculated by subtracting and dividing each trials fluorescence *F(x, y, t)* by the frame preceding the stimulus delivery:

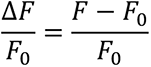

where F_0_(x, y) is the frame of stimulus delivery (F_0_ = F at t = 0). Note, an extended analysis was performed with different normalization methods by subtracting and dividing each trials fluorescence F by the fluorescence averaged across different time windows before stimulus onset (t = 0 ms, t from [-100,0] ms, t from [-200,0] ms). Increasing normalization windows slightly altered the change in fluorescence magnitude and variance of the evoked response; however, varying the normalization window did not affect the adaptive cortical response reported in this study (Suppl. Fig. 3).

A single region of interest (ROI) was identified using the largest 10 x 10 pixel (434 x 434 µm) area response 20-25 ms following stimulus onset. The average activity within this region was extracted across all frames to compute the temporal dynamics of the fluorescent signal. Note, due to the fluorophore^17^, positive changes in membrane potential correspond to a decrease in ArcLight fluorescent activity. In line with our previous study^16^ all traces have been inverted to show a positive increase in fluorescence. Fluorescent voltage traces and behavioral lick responses were acquired within the same time window and aligned with regard to stimulus onset (Fig. 1f). After an animal was lesioned, GEVI responses were qualitatively compared to the responses recorded before lesion. Temporal dynamics and overall fluorescence were assessed.

### Ideal observer analysis

To quantify the fluorescence signal over the course of learning a metric *d*′_*neuro*_ was computed. For a given day or session, single trial distributions of evoked signal peaks (maximum % ΔF/F_0_ within 100 ms post stimulus) were compared to the corresponding noise distributions when no stimulus was present. *d*′_*neuro*_ is then defined as:

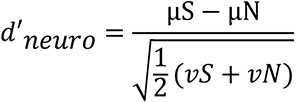

where µS and µN are the mean and *vS* and *vN* are the variance of the signal and noise distribution. *d*′_*neuro*_ was then directly compared with the behavioral learning curve derived from *d*′_*behav*_. The same analysis was repeated for the data acquired with changing stimulus statistics, with the exception that only trials with an intermediate stimulus amplitude shared between the high and the low range condition were used. Note, the chosen stimulus of 8-degree whisker angle represents the midpoint of the high range and the upper limit of the low range condition. Distributions consisting of all evoked trials termed “signal” (peak ΔF/F_0_ with 8 °) and catch trials termed “noise” peak (ΔF/F_0_ with 0 °) were computed separately for the high and low range condition with one switch from high to low or vice versa. Statistical differences between distributions were assessed using bootstrapped estimates of 95 % confidence limits for the *d*′ metrics. This means resampling the *d*′ from a given session with replacement 1000 times, taking the average of each resampled dataset, and then taking the interval that spans the central 95 % of this distribution of averages across resampled datasets. Significance values were further estimated with a non-parametric Wilcoxon rank-sum or Kruskal-Wallis test. Throughout this manuscript, * indicates p < 0.05; ** indicates p < 0.01; *** indicates p < 0.001; and “n.s.” indicates “not significant”.

### Cascade framework

This analytical framework was created to describe the correspondence between the sensory input distribution, the neuronal response function derived from the GEVI signal in S1, and the behavioral readout. The experimentally controlled stimulus amplitude on a given trial *s_i_* is drawn randomly from the input distribution. The evoked GEVI signal in S1 can then be expressed as a stimulus response function G(*s_i_*). To establish a link between G(*s_i_*) and the behavior, a mathematical function *f*(.) was created that transforms G(*s_i_*) to a probability of a lick response *P*(*GO*|*s_i_*), i.e. the psychometric function, as illustrated in Figure 5a and b. In other words f(.) represents a matching of the evoked fluorescence to the lick response, given a particular stimulus. Combining both G(*s_i_*) and *f*(.) results in a function *f*[G(*s_i_*)], as an estimate of the psychometric curve, *P*(*GO*|*s_i_*), that can then be directly compared to the actual psychometric curve *P*(*GO*|*s_i_*). This approach can be applied across the different stimulus conditions (high versus low), and comparisons made across the corresponding G(*s_i_*) and *f*(.) functions.

To quantitatively estimate how much of the behavioral variation can be explained by S1 activity versus downstream, the following control was performed: G(*s_i_*) is considered to change between the high and the low range condition, as observed experimentally, from G*_High_*(*s_i_*) to G*_Low_*(*s_i_*). As a null test for the transition from the high to low condition, to capture how much of the observed changes in behavior is explained solely by the changes in G(*s_i_*), *f*(.) is held constant, operating from a function *f_High_*(.), that only reflects the high range condition. The combination of G*_Low_*(*s_i_*) and *f_High_*(.) then produces an estimated psychometric function 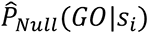 that is only influenced by changes in G(*s_i_*), and thus serves as a null test for the prediction based on changes in neural activity in S1 alone.

Because experiments were designed to present a range of stimulus amplitudes, comparisons of the fluorescence (Fig. 4c) and behavioral (Fig. 5a, b) responses involved comparing curves. One simple way to do this is to compare the area under the curves (AUCs) by taking the difference between the area under the curve across the conditions. For the observed GEVI fluorescence signals for the high and low range stimulus conditions (Fig. 4c), this difference was:

**Figure 4.**
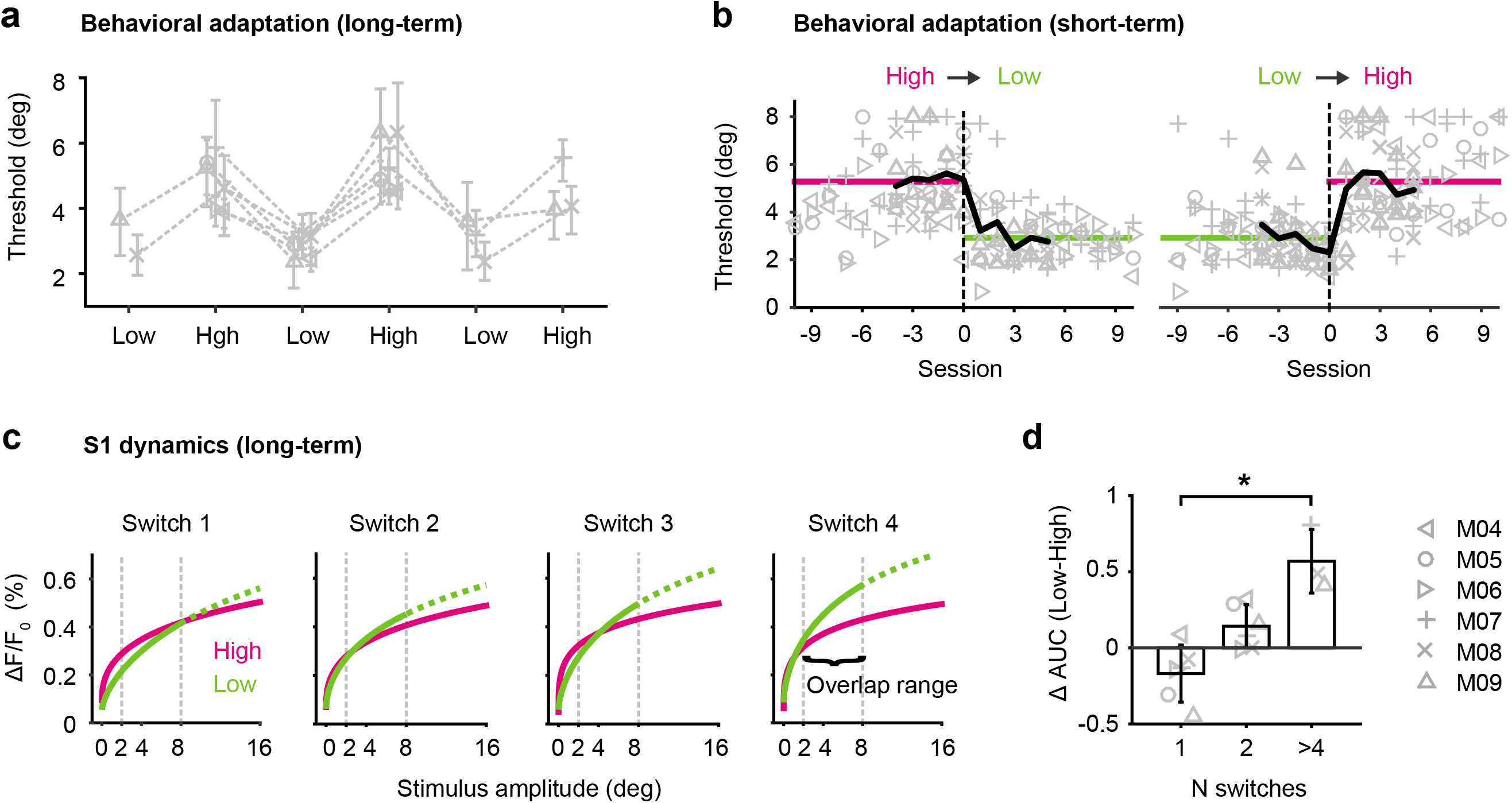
Timescale of behavioral adaptation and related S1 dynamics. **a,** Long-term behavioral adaptation. Psychometric thresholds of individual mice (n=6) undergoing repetitive task changes starting with either the high range or low range condition. Symbols represent mean thresholds across sessions (n=5-11 per condition), errorbars represent 95% confidence limits. Note, conditions are changed in an alternating fashion to test reversibility of adaptation. **b.** Short-term behavioral adaptation at the transition (dashed lines) from high to low (left) and low to high (right). Symbols represent thresholds from each mouse and each session. Bold lines represent thresholds averaged across mice before and after the transition. Horizontal lines represent thresholds averaged across mice and sessions for a given condition. Note, the data is normalized with regard to the transition, with session 0 representing the last session of the preceding condition. **c,** S1 dynamics over multiple task changes (long-term). Neuronal Stimulus response curves derived from the ΔF⁄F_0_ measurement in S1. The curves are weibull fits to the data of n=3 mice undegoing repetitive changes. The “overlap range” is defined as the stimulus range common to both high range and low range datasets (2-8 degree) which is used to calculate the area under the curve (AUC). The hypothetical part (above 8 degree for the low range) is represented as dashed curve. **d,** Difference between high range and low range S1 responses, calculated as the difference in AUC space (AUC_Low_-AUC_High_) at different training stages. Bars represent means across mice with SD (not all mice underwent all switches; switch 1-2, n=6 mice, switch >4, n=3 mice). * P < 0.05; ** P < 0.01; *** P < 0.001; ‘n.s.’ not significant, Kruskal-Wallis test.

**Figure 5.**
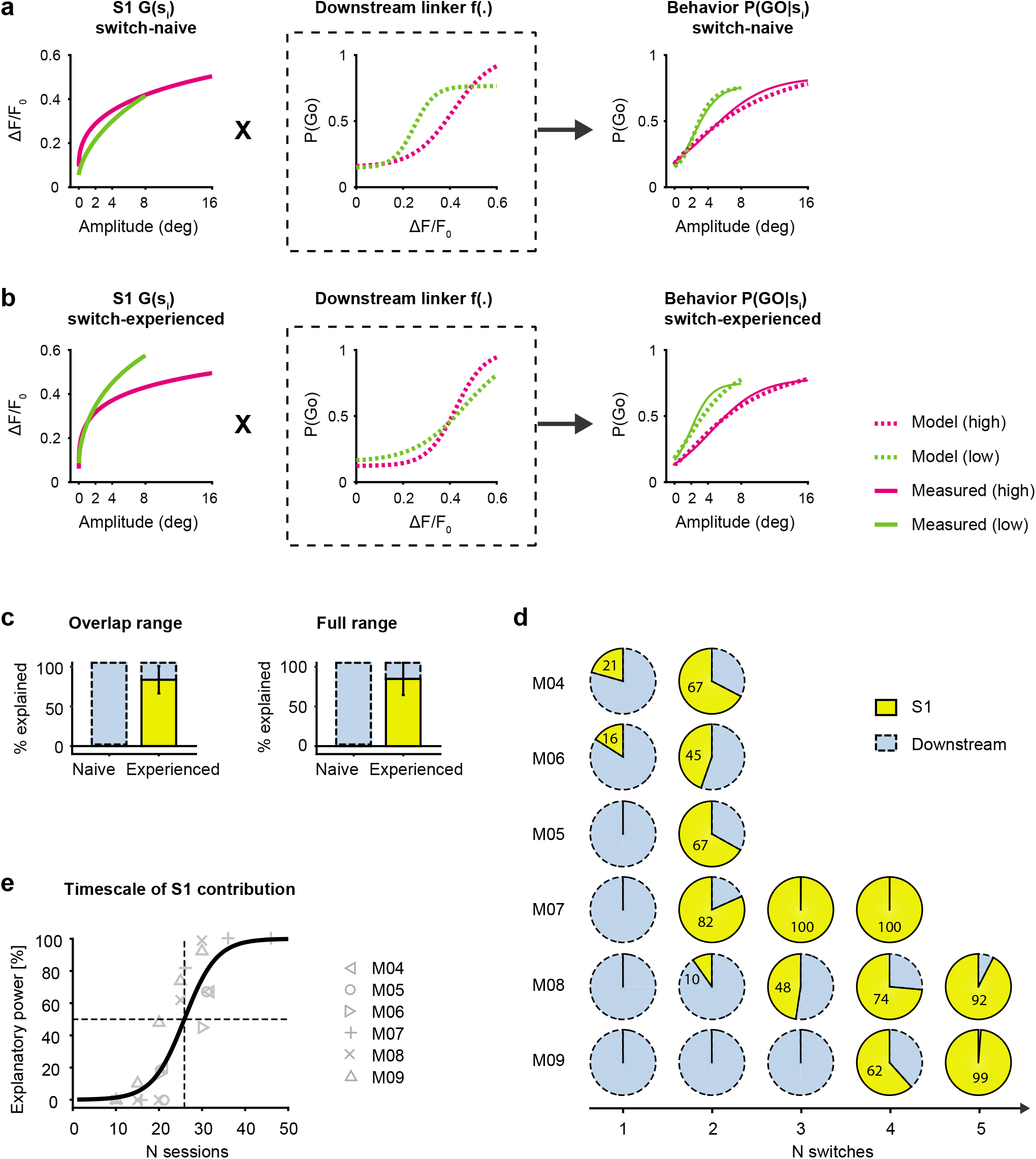
S1 contribution to long-term behavioral adaptation. **a,** Model incorporating the sensory input distribution S_i_, the neuronal response function G(s_i_) derived from the GEVI signal in S1 and the behavioral readout P(GO|s_i_). The function f(.) represents a downstream process, matching the evoked fluorescence to the lick response. Model predictions and measured data at the “switch-naive” training stage, when subjects are challenged with a change in stimulus satistics the first time. **b.** Model predictions and measured data at the “switch-experienced” training stage, when subjects have undergone at least 4 switches. **c,** Relative explanatory power in S1 of mice undergoing multiple changes. Model results are shown separately for the overlap range (left) and the full range (right). Bars represent means across mice with SD (n=3). **d,** Pie plots depict the fraction explained by S1 versus downstream quantified for the behavior of mice undergoing different number of switches and different number of sessions per condition (M04-M06, n=2 switches, n=10-11 sessions, M07-M09, n=4-5 switches, n=5 sessions). **e,** Timescale of S1 explanatory power. The fraction explained by S1 is plotted as a function of the number of sessions. Solid curve is a logistic fit to all data.

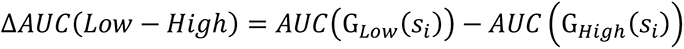

where the area was computed across the “overlap range” of stimulus amplitudes.

Similarly, to quantify the relative explanatory power of S1 versus downstream processing (Fig. 5c, d, e), we implemented the following. The behavioral adaptation was quantified by subtracting the areas of the model fits for the high and low range conditions:

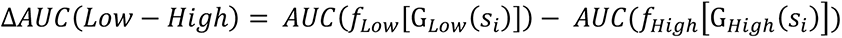

Examples of data and model fits are in the right column of Figure 5a and b. Since, by construction, this analysis involved components from both S1 and downstream of S1, this allowed us to evaluate the relative contributions of each. Assuming that G(*s_i_*) changed from G*_High_*(*s_i_*) to G*_Low_*,(*s_i_*) but *f*(.) is constant (*f_High_*(.)), this resulted in a model prediction 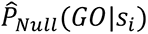 for the behavior performance, if the effect were due to S1 alone. The prediction for the behavioral adaptation based on a change in G(*s_i_*) (and thus S1) alone was then calculated as follows:

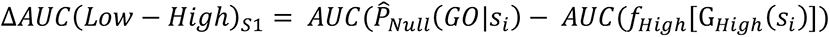

Finally, the fraction explained by S1 was then calculated from Δ*AUC*(*Low* − *High*) and Δ*AUC*(*Low* − *High*)*_S_*_1_ as follows:

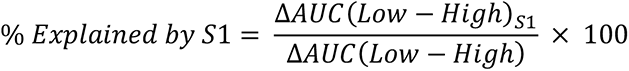

and the remaining was explained downstream of S1, or % *Explained downstream* = 100 −% *Explained by S*1.

All curves were fit using the psignifit toolbox^27^ and the goodness of fit was assessed by calculating metrics of deviance (D) as well as the corresponding cumulative probability distribution (CPE). To rule out the possibility of poor fitting in the cascade framework, we inspected the goodness-of-fit metric of deviance (D) as well as estimates of where the goodness-of-fit lay in bootstrapped cumulative probability distributions of this error metric (CPE) using the psignifit toolbox. Due to the steep increase in ΔF/F_0_ at lower stimulus amplitudes we find that a Weibull function provides the best fit for G(s*_i_*). Both, the linker function f(.) and the psychometric function are best fit by a logistic function due to the sigmoid configuration of the data.

## RESULTS

The current study investigates learning and experience dependent adaptation in the mouse vibrissa system. Our main goal is to establish a relationship between controlled whisker inputs (stimulus), S1 activity, and licking response (behavioral output). We manipulated stimulus inputs and observed S1 activity during early basic detection learning. We then observed these animals longitudinally for flexible adaptation in a changing environment. To repeatedly measure signals of large neuronal pools at different training stages, we performed chronic wide-field imaging of S1 activity with the genetically encoded voltage indicator (GEVI) “ArcLight” in behaving mice.

Figure 1 outlines the experimental design and summarizes the basic neuronal and behavioral metrics of this study. Seven mice were transfected with the GEVI ArcLight before being imaged through the skull using a wide-field fluorescence microscope (Fig. 1a). We have previously described this imaging technique in detail^16^. Figure 1b shows the characteristic spread of ArcLight expression in an example coronal brain section. The same hemisphere is shown *in vivo* in Figure 1c with spontaneous fluorescence activity in S1 at the beginning of a behavioral session. Figure 1d shows sequential frames of typical fluorescence activity patterns in a trained mouse recorded in response to deflections of a single whisker.

Transfected mice were first trained on a tactile Go/No-Go detection task ^22, 23, 26, 30, 31^. Animals detect pulse-shaped deflections of a single whisker and generate a lick on a waterspout (“Go”) or, if no stimulus is present, animals withhold licking (“No-Go”) (Fig. 1e). In response to stimulation, trained mice first show a clear cortical fluorescence response within 100 ms, then stereotypical lick response approximately 200 ms later (Fig. 1f). Note, the temporal resolution of ArcLight allows us to identify the fluorescence response peak in S1 (maximum % ΔF/F0) within the 100 ms window post stimulus which clearly precedes the typical behavioral lick response. Hence, the sensory signal in S1 can be disentangled from potentially interfering motor related signals caused by licking. For early “basic learning” experiments the neuronal S1 signal was observed in mice acquiring the basic principles of the Go/No-Go task, as demonstrated by learned lick responses (Fig. 1g, left). For later “adaptive behavior” experiments (Fig. 1g, right), S1 was imaged in mice challenged with randomly presented whisker deflection amplitudes and systematically manipulated statistical distributions.

### S1 responses during basic learning

All subjects received uncued single whisker stimulations at random intervals of 4-10 s with a single pulse or no stimulation (catch trial). Learning was measured by calculating the hit *p*(*hit*) and false alarm rate *p*(*fa*) of successive daily trainings. A criterion of *p*(*hit*) = 0.95 and *p*(*fa*) = 0.25 was used to determine successful acquisition of the task. Figure 2a shows hit and false alarm rates for three mice trained on the weak stimulus (A = 4°). With this weak stimulus, subjects required extended training for more than 30 sessions to achieve successful performance.

Figure 2b shows frames of cortical S1 fluorescence activity from an example mouse over the course of learning. Each square represents a top view of the same cortical hemisphere on selected training days with the average fluorescent activity (ΔF/F_0_ in %) at the frame corresponding to the peak of the sensory evoked signal (A = 4°, top row) or during catch trials (A = 0°, bottom row). The magnitude and spread of fluorescence is highly variable throughout the “basic learning” daily sessions, and a significant trend cannot be identified with learning progress.

To further quantitatively evaluate the fluorescence signal over the course of learning, we used classical signal detection theory^32, 33^ with a computed neuromeric sensitivity measure d’_neuro_. Single trial distributions of evoked signal peaks were compared to the corresponding noise distributions, when no stimulus is present. The d’_neuro_ for a given day or session was then calculated by subtracting the means of the distributions and dividing by the variance. To compare this metric to the behavior, a d’_behav_ was calculated from the observed behavioral hit and false alarm rate of each training session (see methods). Figure 2c shows the d’_neuro_ (orange) along with the behavioral learning curve d’_behav_ (gray), across all sessions and mice. The dashed lines separate performance into “detect-naive” (d’<0.8), and “detect-experienced” (d’>1.5). In contrast to the behavior, the average neuromeric sensitivity remains at a relatively constant level, representing a stable signal to noise relationship independent of the continuing learning progress. Note, a second group of animals (n = 3) was trained on a much stronger stimulus for comparison (A = 16°). Those mice achieved much higher hit rates at the beginning of training and reached successful task acquisition in half the time (Suppl. Fig. 1a); however, the neuromeric sensitivity was also orthogonal to the learning progress (Suppl. Fig. 1b-c). We conclude that neuronal sensitivity in S1 measured from a large population of neurons does not change during basic learning.

### Adaptive behavior and changes in S1

To investigate behavioral and neuronal dynamics with regard to changing context, we performed experiments in which we systematically manipulated stimulus statistics (Fig. 3a). The psychophysical techniques were adapted from a behavioral paradigm we recently developed in the rat^26^ which are described in materials and methods. The first distribution consists of four different stimulus amplitudes of whisker deflection, as well a catch trial (A = [0, 2, 4, 8, 16] °, magenta) which we refer to as the “high range” condition. The second distribution consists of four new stimulus amplitudes of whisker deflection, and a catch trial (A = [0, 1, 2, 4, 8] °, green), which we refer to as the “low range” condition. Each stimulus or catch trial was presented with equal probability (uniform distribution). Importantly, the experimental design involves amplitudes common to both high-range and low-range conditions, three of the four stimulus amplitudes were shared between conditions. Figure 3b depicts typical psychometric curves from an example mouse, performing the task first under the high range (magenta) and low range condition (green). There is a consistent shift of the psychometric curve in response to the changing stimulus statistics, which we refer to here as “adaptive behavior”. All animals experienced both a switch from high range to low range, as well as low range to high range. Adaptation was reversible, meaning that over multiple switches, the animal’s performance changed, indicating that they are able to apply a reciprocal task strategy, modulated by the shift in stimulus distribution.

The simple reward expectation model tests if the animal adapts its behavior to maintain accumulated reward in the face of a changing stimulus distribution. If the animal does not adjust its behavior the predicted low range psychometric function would align with the high range psychometric function (Fig. 3b black dotted curve under magenta curve; H_0_). In moving from the high range to the low range condition, this would result in a decreased reward rate for the same number of trials. However, if the animal adjusts its behavior, the model predicts that the psychometric function shifts to the left such that the expected reward per trial remains constant (Fig. 3b black dashed curve; H_1_).

We observed a shift of the psychometric curve when going from high range to low range performance. The experimentally measured psychometric function in the low range condition (green) comes quite close to the hypothetical performance, suggesting that the animal adapts its behavior to maintain reward. The psychometric shift results in a significant decrease in response threshold for all mice (Fig. 3b inset). Figure 3c depicts the actual trial-by-trial reward accumulation by the example mouse. Overlaid are results for n = 11 sessions with the high range distribution (magenta) and n = 10 sessions with the low range distribution (green). The slope of reward accumulation in the low range condition nearly matches that of the high range condition, and the slope for the low case (green) is close to the prediction from the maintenance of accumulated reward hypothesis, H1 (dashed line), while being clearly separable from the slope representing the null hypothesis (dotted line). The total number of rewards acquired on average per session and across all mice further confirms this (total # high range = 44.5 ± 8.6, total # low range = 42.0 ± 10.6, Mean and SD, Fig. 3c inset), whereas there was no evidence for an alternative strategy to maintain the total number of rewards by working substantially more trials. The total number of trials per session was nearly the same for each condition on average (high range, n = 101 and low range, n = 104, Fig. 3c inset). These findings show that detection behavior is highly flexible in the face of a changing stimulus distribution.

Figure 3d shows frames of evoked S1 fluorescent activity in a mouse exposed to high range stimuli (top) or low range stimuli (bottom). In both conditions, we observed an increase in fluorescence amplitude and spatial spread, with increasing stimulus amplitude. Note, there is a relatively consistent linear relationship between the percent change in fluorescence magnitude and the activated cortical area, as both scale with stimulus strength in a highly correlated fashion (Suppl. Fig. 2). For simplicity, we use percent change in fluorescence magnitude as a metric for cortical activation. The evoked activity for the same stimulus is clearly higher in the low range compared to the high range condition.

To further dissect this change, we focused our analysis on trials with an intermediate stimulus shared between both datasets (outlined box in Fig. 3d). The chosen whisker deflection amplitude of 8 degrees represents the midpoint of the high range distribution and the upper limit of the low range distribution. Figure 3e depicts the temporal fluorescent signal in response to an 8° amplitude, averaged across sessions from all four mice. These mice have been exposed to both high and low range conditions as well as a switch between conditions. Histograms of behavioral lick responses are shown on top – the lick response occurs approximately 200 ms window after the measured fluorescence peak. We did not observe a significant difference in behavioral response latencies between the high-range and the low range condition. However, the evoked S1 response to the same 8° deflection was higher in animals challenged with the low range vs the high range stimuli.

Figure 3f outlines the observer model we used based on signal detection theory, to interpret and analyze data from the S1 neural signals, as well as a presumed downstream decision criterion. Multiple scenarios are considered to trigger a change within sensory and higher order processing stages; behavioral adaptation can be induced by changes in sensitivity (d’) intrinsic to S1, changes in decision criterion (c) by a downstream observer, or both. Sensitivity in S1 improves through reduction in the overlap between sensory signal and noise distributions (top panel). In addition, the downstream observer may value hits and false alarms differently by altering the decision criterion (bottom panel). To test these predictions, distributions consisting of many evoked trials termed “signal” (ΔF/F_0_ with 8 °) and catch trials termed “noise” (ΔF/F_0_ with 0 °) were computed separately for the high and low range condition (Fig. 3g, top). The noise distribution (grey) is comparable for the high and the low range condition with its means being identical (dashed vertical black lines). However, the signal distribution for the low range (green) is shifted towards higher fluorescence changes as compared to the signal distribution of the high range (magenta) with a clear difference in its mean (dashed vertical magenta and green lines). As shown in the bottom of Figure 3g, a significant difference in neuronal sensitivity was confirmed by calculating the neuromeric d’ as introduced earlier in this study. While the neuronal d’ was altered slightly by different normalization methods of the ΔF/F_0_ metric, an extended analysis revealed that the difference in d’ across conditions persisted (Suppl. Fig. 3a-c). In addition, we created receiver operating characteristic (ROC) curves by varying a criterion threshold across the ΔF/F_0_ signal and noise distributions and plotting the hit rate (signal detected) against the false alarm rate (incorrect guess) (Suppl. Fig. 3d). The ROC curve for the low range condition was higher than that for the high range condition, quantified by a larger area under the low range ROC curve (AUC_low_ = 0.77) than for the ROC curve for the high range condition (AUC_high_ = 0.69), thereby confirming a change in S1 sensitivity.

Changes in criterion were inferred by comparing the hit rate in ROC space (neurometric) with the average hit rate measured from the behavior (psychometric). The criterion shows a slight, yet non-significant decrease when operating from the low range compared to the high range condition (Fig. 3g, bottom left, blue). Note, the S1 sensitivity and downstream criterion change in opposite direction, e.g. an increase in hit rate can be caused by an increase in S1 sensitivity and/or a decrease in criterion. We conclude that, in highly trained animals there is evidence of adaptive sensitivity in S1, yet comparatively smaller adaptive changes in criterion by a downstream observer.

### Changes across training stages

The data described thus far, comparing the adaptive behavior and cortical activity across the high and low range conditions was derived from highly trained subjects. To better understand the development of this phenomenon over time we re-analyzed the data from highly trained subjects at different training stages. Conditions were always changed in alternating fashion to test the reversibility of adaptation (high-low-high etc.). Figure 4a shows the psychometric threshold for mice undergoing repetitive task changes starting with either the high range or low range condition. We observed that the order of stimulus conditions does not affect behavioral adaptation and response thresholds closely follow the direction of task manipulation. This suggests that mice are able to apply a flexible and reciprocal task strategy modulated by the shift in stimulus distribution. Behavioral adaptation was stable across multiple training stages, as revealed by consistently modulated thresholds.

Since individual mice experienced different numbers of switches (n = 2-5) and different numbers of sessions for a given condition (n = 5-11 per condition), we asked how many sessions or trials are generally required to adapt after a switch? We focused our analysis on the transition from one condition to another. Figure 4b shows psychometric thresholds extracted from consecutive sessions at the high- to low range transition (left) and at the low- to high range transition (right). The data were normalized with regard to the transition and data from all switches were included in this analysis. Across mice, the target threshold for the high or low range condition (average threshold per condition) was reached within approximately three sessions, indicating that behavioral adaptation to changing stimulus statistics required repeated exposure to approximately 300 trials within a given condition.

In order to investigate S1 dynamics during long-term adaptive behavior, GEVI data were acquired and analyzed across different training stages. Figure 4c shows neuronal stimulus response as a function of stimulus amplitude derived from the ΔF/F_0_ measurement in S1 of three mice that were challenged with a minimum of four switches. The functions are represented by a curve fitted to the average data (see methods), and separately shown for each high range and low range condition. Comparing the S1 response curves across different training stages revealed an interesting pattern; at the first switch, high- and low range curves are similar despite differences in the corresponding stimulus distributions. However, high- and low range curves progressively separate as a function of switches and training experience. Since the fitting procedure is constrained by data collected across different stimulus ranges, we focused our analysis on the “overlap” range which is defined as the stimulus range common to both high- and low range datasets (A = [2, 4, 8] degrees, dashed vertical grey lines). This range was then used to calculate the area under the curve (AUC) for each condition, as a way to compare the two curves (see methods). Figure 4d shows the difference between high- and low range S1 responses, calculated as the difference in AUCs of the low and high range response curves (Δ*AUC* (*low* − *high*)). S1 responses adapt progressively and systematically as a function of training experience. To determine whether this neuronal effect was dependent upon the animals’ choice, we conducted a separate analysis by tracking the stimulus evoked S1 signal across different training stages, separately for HIT and MISS trials (Suppl. Fig. 4). The results from the parsed trials for either HITs or MISSes (Suppl. Fig. 4a-b) are very similar to the combined trials (Suppl. Fig. 4c), confirming that the emerging difference in S1 activity is not biased by the choice element.

### S1 contribution to adaptive behavior

The core elements offered through the behavioral and neurometric measures analyzed thus far provide somewhat limited “slices” of the underlying phenomena. We thus sought to develop a more comprehensive analytic, cascade framework that enables us to evaluate the relative contributions from activity in S1 and downstream of S1 in predicting the observed behavior across the full range of stimuli. The framework establishes a link between the sensory input distribution s_i_, the S1 response function G(s_i_), and the psychometric curve P(GO|s_i_) (Fig. 5). The function f(.) represents a downstream process, matching the evoked fluorescence to the lick response, given a particular stimulus, and is referred to as the “linker” function, as it links S1 activity to behavior. This framework can then be used to describe the changes underlying the adaptive behavior and assess the relative roles of the two stages, S1 and downstream (see methods for details). Animals that were challenged with a single switch were defined as “switch-naive”; animals that were challenged with a minimum of 4 switches were defined as “switch-experienced”. Figure 5a and b show the different elements of the cascade framework applied to data from multiple training stages, allowing us to separately investigate switch-naive (Fig. 5a) and switch-experienced (Fig. 5b) transformations.

S1 response curves are shown separately for the high- and low range conditions (Fig. 5 a, b; left panels), revealing a strong separation at the switch-experienced level. In contrast, the behavior data (Fig. 5 a, b; right panels) are consistently modulated by changes in the stimulus distribution at both training stages. This asymmetric relationship of S1 activity and the behavioral output has an impact on the downstream linker function (Fig. 5a, b; middle panel). At the switch-naive training stage (Fig. 5a; middle panel), high and low range linker functions appear different, indicating that the associated neuronal processes that are predictive of behavior, are not S1 based. In contrast, at the switch-experienced stage (Fig. 5b; middle panel) high- and low range linker functions are relatively congruent, transforming the modulated S1 response functions directly into an adapted behavioral response. All predictions from combining S1 response functions and linker functions (Fig. 5a, b; right panel, dashed curves) match well with the measured psychometric curves (Fig. 5a, b; right panel, solid curves), with minor differences due to the fitting accuracy when generating both S1 and linker functions. To capture how much of the observed changes in behavior is explained solely by changes in S1, we used a null test approach assuming a fixed linker function (e.g. high range only) and predicting the change in psychometric curve (e.g. low range) based only on changes in S1 response (e.g. from high to low range). The behavioral adaptation was estimated by subtracting the areas under the curve of the low- and high range psychometric model predictions, again as a way of comparing the two curves, and the fraction explained by S1 was then estimated by comparing this difference with the null test results (see methods for details). Figure 5c shows the relative explanatory power summarized for mice at the switch-naive versus switch-experienced training stages and separately for the overlap range (left) and full range (right). The results of both analyses reveal that changes in S1 activity cannot explain behavioral adaptation at the switch-naive stage (0 %), whereas they can explain a large proportion (85 %) at the switch-experienced level. The primary behavioral adaptation occurs in regions outside of S1. However, as animals gain experience in a changing sensory environment, adaptive responses emerge in S1.

In addition to a changing environment, switch-experience also incorporates increased exposure to both stimulus conditions and can be described as session-experience. We applied the cascade model to each animal individually to test for the effect of session experience vs switch experience. Figure 5d depicts the relative explanatory power of S1 versus downstream, quantified for all mice at each transition. The first group of mice (M04-M06) experienced only two switches but a higher number of sessions within a condition (n = 10 sessions), whereas the second group (M07-M09) experienced four to five switches but a smaller number of sessions within a condition (n = 5 sessions). Despite the variability across mice, the S1 explanatory power increased collectively with training, again suggesting that the adaptive changes are increasingly emerging in S1 with experience to the point where a large percentage of the behavioral changes are reflected in the sensory evoked S1 responses. Figure 5e presents the explanatory power of S1 as a function of the number of sessions across animals, showing a progression across sessions, reaching 50 % explanatory power between 20-30 sessions. Interestingly, mice challenged with a smaller number of sessions per block required more switches for the explanatory power to emerge from S1, suggesting the number of switches and number of sessions, are both important determinants of this experience dependent effect.

To causally test whether context-dependent representation in S1 ultimately contributes to behavioral adaptation, we chronically inactivated S1 in a subgroup of trained mice with the neurotoxin ibotenic acid, where GEVI activity was used to facilitate targeted acid injections (Fig. 6). Both, simple detection and adaptation to changing conditions were tested in healthy (n = 6), lesioned (n = 4), and sham lesioned (n = 2) mice, and compared either within the same group or across groups. From the four lesioned mice, two were switch-naive and two were switch-experienced. Figure 6a shows a wide-field image of a coronal mouse brain section (100 μm, right hemisphere) with cytochrome oxidase staining, showing an example S1 lesion. Figure 6b shows frames of cortical fluorescence activity from an example mouse working on the detection task pre- and post-lesion with a clear absence of evoked cortical fluorescence after the lesion (bottom row). Interestingly, mice could still detect stimuli quite well, reaching a level of performance comparable to pre-lesion levels after one day of recovery, as indicated by the psychometric threshold (Fig. 6c). These results agree with recent work showing that S1 is not required for simple detection^12^. However, if challenged with changing stimulus conditions (high-versus low range), the adaptive behavior was clearly compromised following the lesion as shown by the psychometric curves and corresponding psychometric thresholds in Figure 6d. We observed a significant decrease in switch-behavioral adaptation (Δ threshold) in all animals (Fig. 6e), demonstrating that S1 is necessary for behavioral adaptation. There were no differences in animals that were lesioned at the “switch-naive” stage versus “switch-experienced” stage.

**Figure 6.**
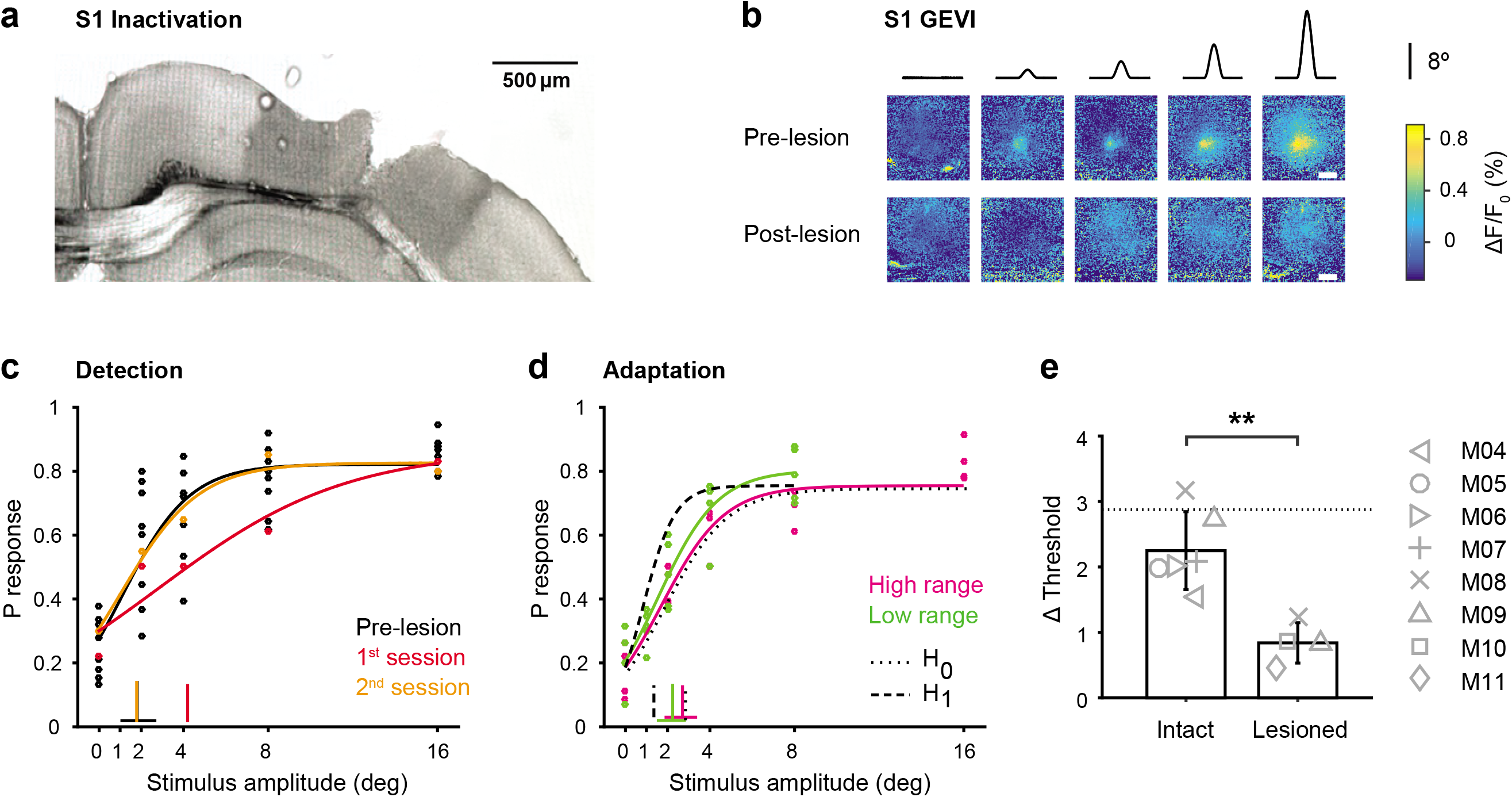
The effect of S1 inactivation on behavior. **a,** Wide-field image of a coronal mouse brain section (100 μm, right hemisphere) with cytochrome oxidase staining, showing a S1 lesion upon acid injection (0.3 μL ibotenic acid 10 mg/mL). **b,** Frames of cortical fluorescence activity from an example mouse before and after the lesion. Top. Example session with S1 intact, showing GEVI activity patterns in response to different stimulus amplitudes (high range, n=25 trials/stimulus). Bottom. Example session of the same mouse after the lesion (n=18 trials/stimulus). Note, the GEVI activity pattern was used for targeted acid linjections. **c,** Psychometric curves and response thresholds for an example mouse detecting high range stimuli before and after the lesion. Each dot corresponds to response probabilities from a single session. Solid curves are logistic fits. Note, pre-lesion data is shown for several sessions (n=5) whereas the post-lesion data is shown seperately for the first (red) and second session (orange) upon lesion. **d,** Psychometric curves and response thresholds for the same mouse challenged with a change in stimulus statistics (high range to low range) upon S1 inactivation. Solid curves are logistic fits to the average data (n=4-5 sessions). Otherwise, figure conventions are the same as in Figure 3. **e,** Behavioral adaptation is calculated as the change in psychometric threshold (Δ threshold) for healthy and lesioned animals. Bars represent means across mice with SD (n=6 intact, n=4 lesioned), dotted line represents mean across sham operated mice (n=2).* P < 0.05; ** P < 0.01; *** P < 0.001; ‘n.s.’ not significant, two-sided Wilcoxon rank-sum test.

Our results suggest cumulatively that the S1 cortical region is not necessary for simple stimulus detection; rather, it plays an important function in experience dependent stimulus adaptation. This highly conserved brain region is likely to be important for an animals’ survival strategies in a dynamically challenging sensory environment.

## DISCUSSION

In this study, we have investigated learning and experience dependent behavior in the mouse somatosensory system. Our findings provide evidence that activity in the highly conserved primary somatosensory cortex can be remarkably dynamic in support of flexible sensory processing and experience dependent behavioral adaptation.

We present the following novel aspects. First, S1 population activity reflects a reliable signal to noise relationship that is driven by the sensory input and does not change during the basic learning process. Second, mice can modify detection behavior in a way as to maintain reward in the face of changing statistical properties of the stimulus. Third, S1 activity is highly dynamic in the face of a changing sensory environment, predicting behavioral adaptation as individuals gain experience. Fourth, S1 is required for this adaptive behavior.

Learning occurs when an individual forms an association based on a new stimulus or context. This process provides obvious benefits such as flexible hunting, optimal foraging, and social communication, especially in environments that tend to change frequently and unpredictably. There is no doubt that associative learning can occur in animals without cortex, including all classes of vertebrates34 and a large number of invertebrate species^35^. Further, several studies elucidating the role of S1 with classical or operant conditioning have found that chronic lesioning of S1 does not affect basic detection12,36, a finding we corroborate here. It is simultaneously the case that other studies demonstrate the opposite effect with acute inactivation of S1^37, 38^, suggesting that S1 is normally central to this behavior. Although this is a complex issue, these studies together seem to suggest that S1 is nominally used for these behaviors, but that given sufficient time for remapping/learning, other pathways are sufficient, and thus S1 may be viewed as involved in but not critical for this simple behavior. A central finding of the current study, however, is that S1 is important for the experience-dependent adaptive behavior *for the exact same task* and does not recover following lesioning of S1, pointing to a critical role for S1 in context dependent, adaptive strategies in a changing environment that could underlie a range of behaviors in more natural settings.

The general behavioral question posed here relates to how the animal responds to changes in the sensory environment. The behavioral paradigm was designed as a highly simplified but carefully controlled manipulation of the statistical distribution of the magnitude of whisker deflections experienced by the animal. Aside from matching amplitudes and velocities of the whisker movement that have been described in a range of studies^20, 21, 39–41^, the current study does not attempt to place this in the context of the natural sensory environment for the animal, and the passive stimulus paradigm does not speak to active sensing that relies on the animal’s own movement in acquiring sensory information. Importantly, although centered around a simple switching in the stimulus statistics, these switches were implemented with multiple changes occurring with alternating conditions, showing that S1 sensitivity closely followed the direction of changes in a way that increasingly explained the behavioral adaptation (Fig. 5), suggesting that the findings here likely generalize to more complex scenarios in the dynamic landscape of the natural sensory environment. The relative role of S1 in the adaptive behavior was found to be strongly experience dependent, where experience is likely to be a complex function of time and exposure to specific aspects of the environment (here number of sessions), as well as the nature, degree, and frequency of change in the environment (here number of switches), which remains to be explored in more detail in naturalistic settings. Overall, the behavior here reveals an active strategy by the animal that robustly maintains reward in the face of a changing environment. The behavior is reversible and independent of the direction of switching, and the animals do not just remain at a higher level of behavioral performance once achieving this or simply work longer sessions to accumulate the same amount of reward. This demonstrates that the phenomenon here is part of an active strategy that involves a cognitive cost to the effort, be it effort on a trial-by-trial basis or the cumulative effort across a session. The changes in strategy we observed were purely reflected in performance, and not in any changes in peripheral sensing such as whisker position or movement given the high degree of control exerted by our passive stimulation paradigm. However, these elements likely play a key role in the overall behavioral strategy in more ethological behaviors. How the behavior, and corresponding sensory representations, change in more complex, naturalistic settings, and at a finer time resolution, are important, and need further investigation in future studies.

While the spatial and temporal resolution of the GEVI imaging here does seem to capture the appropriate scale for the simple detection task, it targets the aggregate activity of large neuronal pools restricted to approximately cortical layer 2/3, and thus does not capture the details of individual neurons of specific cell sub-type across layers that conventional electrophysiological approaches would provide. However, given the long literature implicating the role of inhibitory interneurons in functional plasticity in cortex even on short time scales^42, 43^, we would predict that the adaptive changes in S1 described here would correspond to a differential change in inhibitory drive, and that cortical layer 4 and deeper cortical layers would exhibit less adaptive properties as compared to more superficial layers of cortex, due to the differential dependence upon direct thalamic drive reported across cortical layers^44^. How the behavioral phenomenon we describe here is ultimately driven by population activity across the cortical network is left to more detailed electrophysiological studies in the future. Nevertheless, our finding of context dependent adaptive responses in S1 is surprising as it suggests that reward based choice signals might shift across the cortical network and can appear and influence sensory representation in S1 once an individual has successfully adapted its behavioral strategy. In this context it is important to note that the behavioral adaptation itself is similar at different experience levels, showing consistent changes in performance already before the adaptive response even appears in S1 (switch-naive performer). This is important, as the change in S1 we observe through GEVI imaging averaged across trials is therefore not simply reflecting a difference in performance/choice (e.g. ratio of hits to misses) across the levels of experience, but instead a true change in the S1 response with experience in the context of this task, further reinforced by a direct analysis of the choice signaling (Suppl. Fig. 4). The findings here are thus not in conflict with the previous reports of choice signaling in S1, but instead represent a context dependent element related to more complex behaviors that are likely part of a larger framework of decision making that can only be precisely identified through causal manipulations in future studies.

So, what are the downstream targets that drive early behavior and ultimately change the stimulus representation of primary sensory areas? A recent study found that the perirhinal cortex, the last station in the medial-temporal loop projecting to S1, acts as a gate for the enhancement of corticocortical inputs, which are necessary for basic detection learning^45^. Another recent study probing flexible decision making in the somatosensory pathway of mice found that orbitofrontal cortex dynamically interacts with S1 triggering plasticity based on value signals^15^. In addition, other important reward based choice signals have been reported to influence neuronal signaling throughout cortex^46^. Furthermore, studies of visual attention in primates have distinguished changes in neuronal sensitivity from an observer’s response criterion in extra striate cortex, superior colliculus, as well as lateral prefrontal cortex^47–49^. Hence, there is a large number of downstream candidates potentially modulating the sensory signal stream. Admittedly, in the current study we did not record in any other area outside S1, but the effects described by those other studies collectively point to the interesting idea that the neuronal signal transfer identified by our study could be a common principle across neocortex with “earlier” stages emerging with experience. Moreover, we propose that this shift in cognitive signaling could even affect subcortical structures such as primary thalamus, which we have recently shown to exhibit cognitive signatures in highly trained animals^25^.

Arguably, our current study was not specifically designed to identify attention as a driver for adaptive behavior, even though the measured changes in performance are indicative of a somewhat conscious change in the subjects’ behavioral strategy. Other effects related to the subjects’ thirst, satiation, or arousal level within this behavioral framework can further be excluded, as we have shown previously^26^. However, although speculative, the experience dependent change in S1 sensitivity and the corresponding change in downstream criterion may be signatures of a higher order cognitive process that could be described as an “attentional spotlight” or gain control^50^ moving down the cortical hierarchy. Such a brain-wide dynamic process would facilitate selective and efficient cognitive processing as an individual is adapting to changes in the environment. This transfer could explain some of the disparities in the literature and therefore needs to be further investigated in future studies.

The primary sensory cortex is classically described as the origin of sensory signals in response to stimuli. We have described here a novel role for S1 in long term adaptive behavior. We propose that this highly conserved region has a dual function that is temporally regulated, relaying sensory signals in an acute reactionary fashion vs processing and incorporating dynamic signals into an adaptive behavioral response longitudinally. This suggests primary sensory cortex may not only be involved in fundamental sensory representations, but also long term adaptive strategies which could be necessary for survival.

## Data availability

The data that support the findings of this study are available in “Dryad” with the identifier https://doi.org/10.5061/dryad.h18931zmm.

## Code availability

All computer code are available in “Dryad” with the identifier https://doi.org/10.5061/dryad.h18931zmm.

## Acknowledgements

C. W. was supported by a fellowship from the Deutsche Forschungsgemeinschaft (GZ: WA 3862/1-1) and NIH Brain Grants R01NS104928 and U01NS094302. P.Y.B. was supported by an NIH NRSA pre-doctoral fellowship F31NS09869. M. E. D was supported by NIH Brain Grant R01NS104928. G. B. S. was supported by NIH Brain Grants R01NS104928 and U01NS094302. We thank Cornelius Schwarz and April Reedy for helpful feedback on the manuscript. We also acknowledge April Reedy with the Emory University Integrated Cellular Imaging Core for assistance with histological imaging.

## Author contributions

C.W. and G.B.S conceived the project. C.W., P.Y.B. and M.E.D. designed the study. C.W. and M.E.D. carried out all experiments. C.W., P.Y.B. and M.E.D. analyzed the data, C.W. and G.B.S. wrote the manuscript with comments from P.Y.B. and M.E.D.

---

**Supplemental Figure 1.**
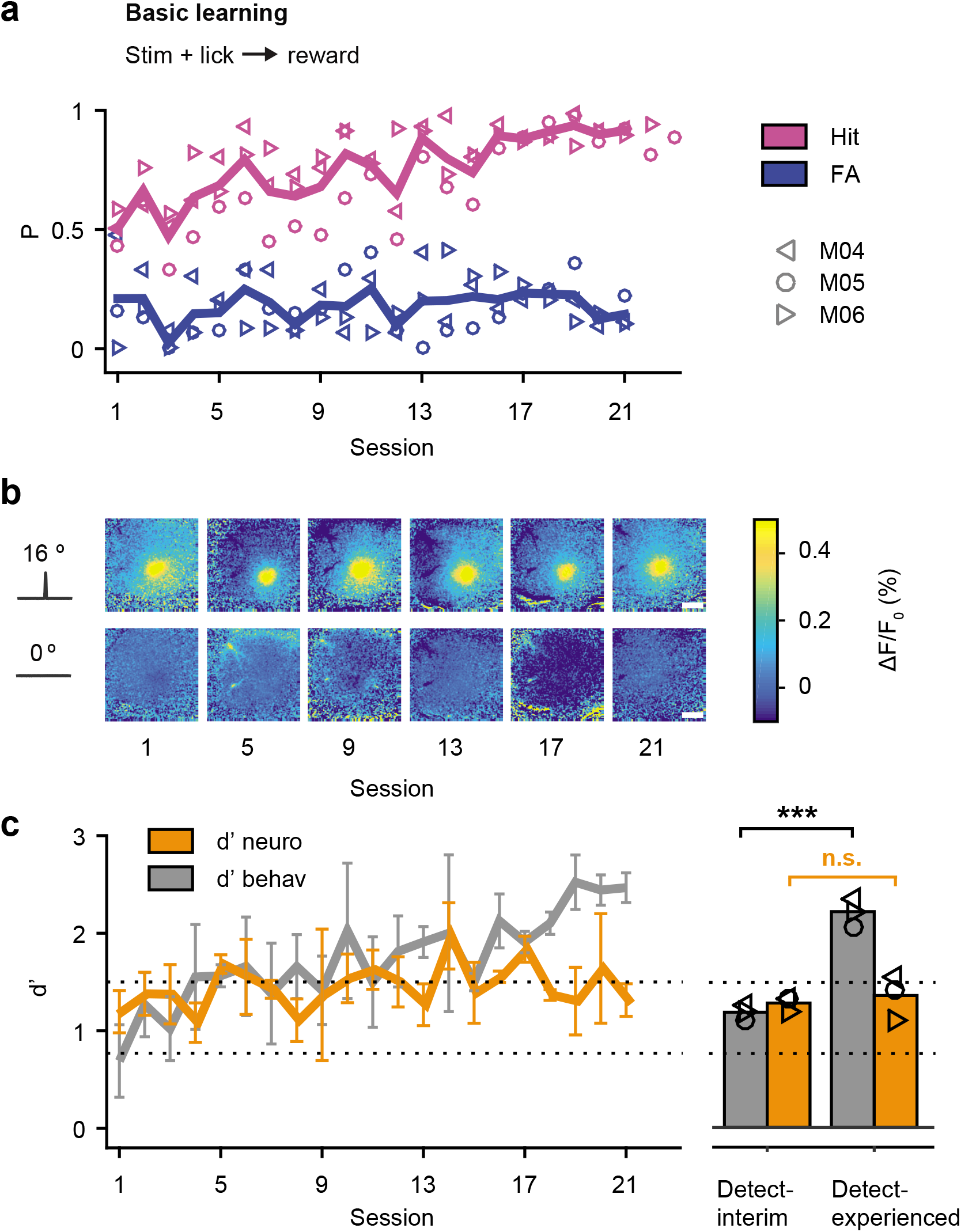
Basic Learning with a strong stimulus. **a,** Learning curve for 3 mice trained on the basic Go/No-Go detection task with a strong stimulus (16 degree). **b,** Fluorescent activity in S1 from an example mouse during learning. Shown are frames at the peak response to a 16 degree stimulus, catch trials are shown below. Scale bar: 1 mm. **c,** Dprime metrics for both behavioral and neuronal data during learning of the task. Note, mice achieved higher hit rates at the beginning of training and reached successful task acquisition in half the time if compared to mice detecting a weak stimulus (Fig 2). The dotted lines separate performance into “detect-interim” (d’=0.8-1.5), and “detect-experienced” (d’>1.5). The right panel shows the same data separated for individuals (symbols) before and after learning. Otherwise, figure conventions are the same as in Figure 2.

**Supplemental Figure 2.**
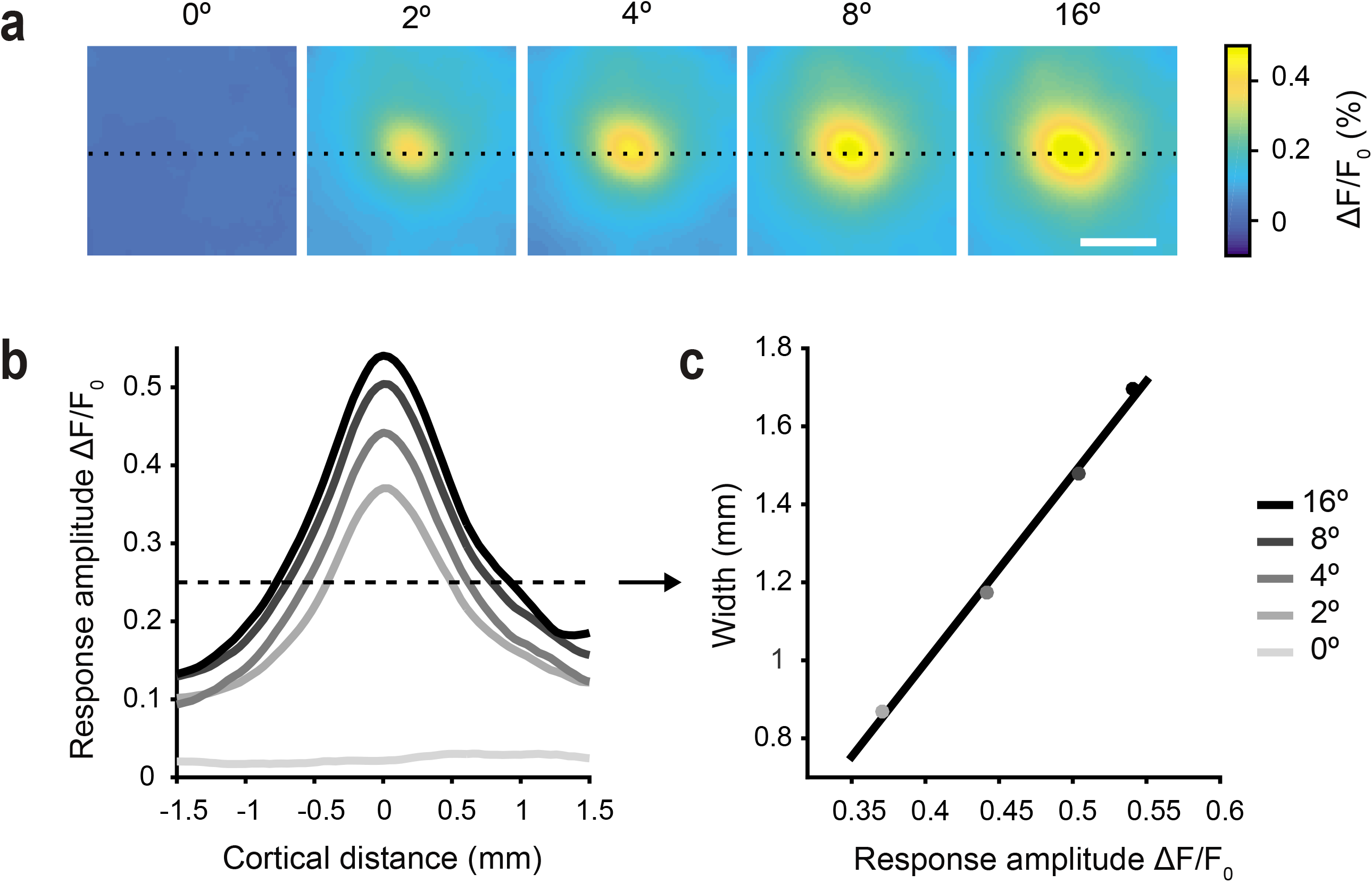
The relationship of magnitude and activated area in GEVI imaging. **a,** Fluorescence activity in response to different stimuli. Each frame is normalized to the frame at stimulus delivery (ΔF⁄F_0_=F-F_0_/F_0_). Shown are images at the response peak. The dotted line represents a slice through the images aligned with the maximum fluorescence. Scale bar: 1 mm. **b,** Magnitude (response amplitude ΔF/F_0_) versus cortical area extracted from the slice in a. Activity patterns are shown in different shades of grey for different stimulus amplitudes. **c,** Relationship between fluorescence magnitude and width of activated area, both scale with stimulus strength in a highly correlated fashion. The width is derived from a threshold (dashed line in b). Data points correspond to different stimulus amplitudes fitted with a linear regression. Shown are means across mice (n=4) and sessions (n=40).

**Supplemental Figure 3.**
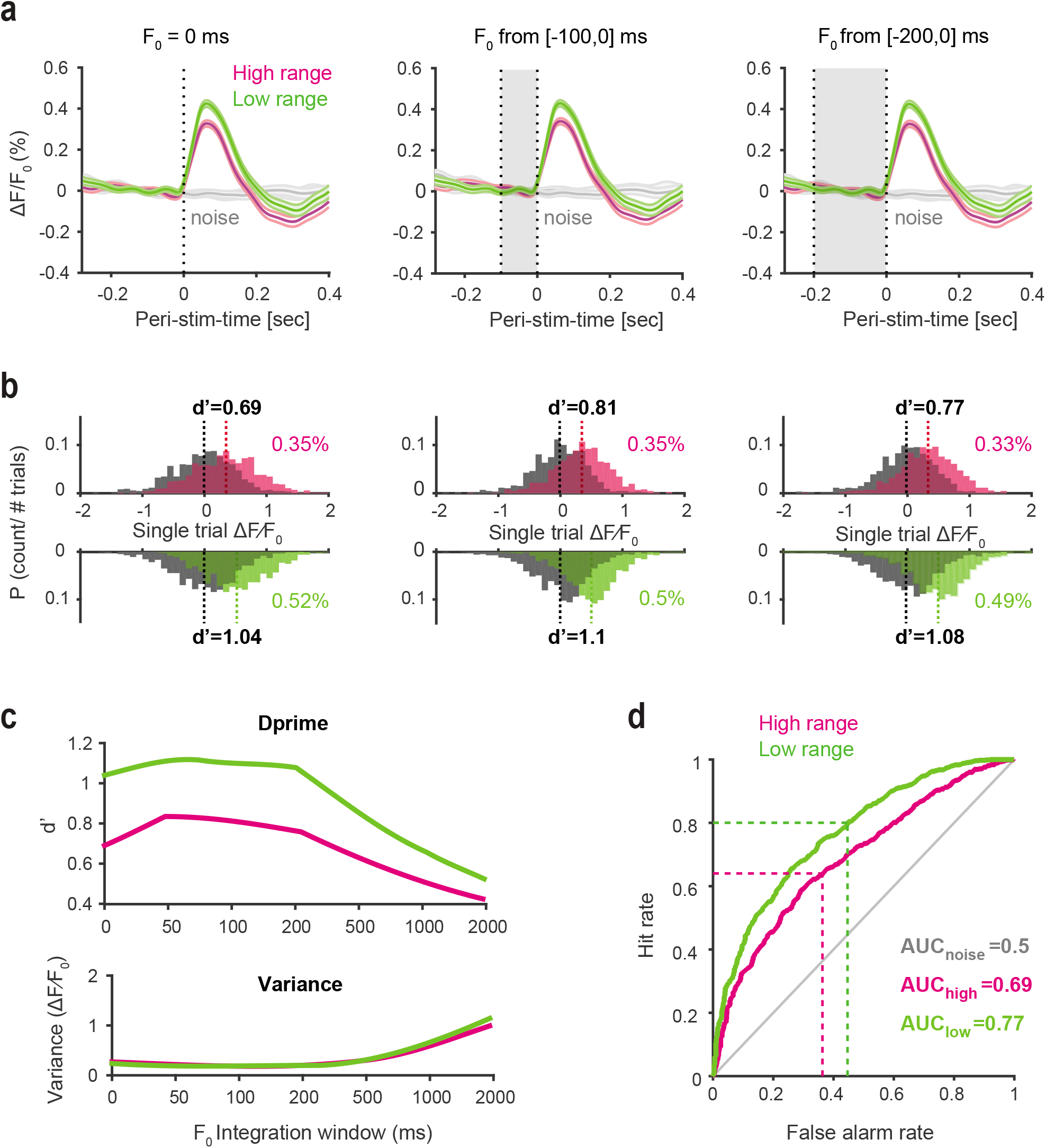
Analysis to test stability of adaptive S1 response. **a,** Different normalization methods. F_0_ was varied by using the mean fluorescence across different time windows before stimulus onset. The ΔF⁄F_0_ measurement was then calculated by F-F_0_/F_0_. Mean fluorescence traces (n=4 mice, 831-870 trials) in response to the high (magenta) and the low range (green) stimulus condition. The grey box depicts the window for calculating F_0_. From left to right: F_0_=0 ms (frame at stimulus delivery), F_0_ from [-100,0] ms, and F_0_ from [-200,0] ms. **b,** Single-trial signal-peak and noise distributions from the data in a. Magenta and green numbers are mean fluorescent values in % ΔF⁄F_0_ for the high and low range condition respectively. Black numbers represent dprime metrics. **c,** Dprime and variance for different F_0_ calculations. **d,** Receiver operating characteristic (ROC) curves created by shifting the criterion across the ΔF/F_0_ signal and noise distributions of the high and low range condition. The downstream criterion can be inferred by comparing the hit rate in ROC space with the average behavioral hit rate (dashed lines).

**Supplemental. Figure 4.**
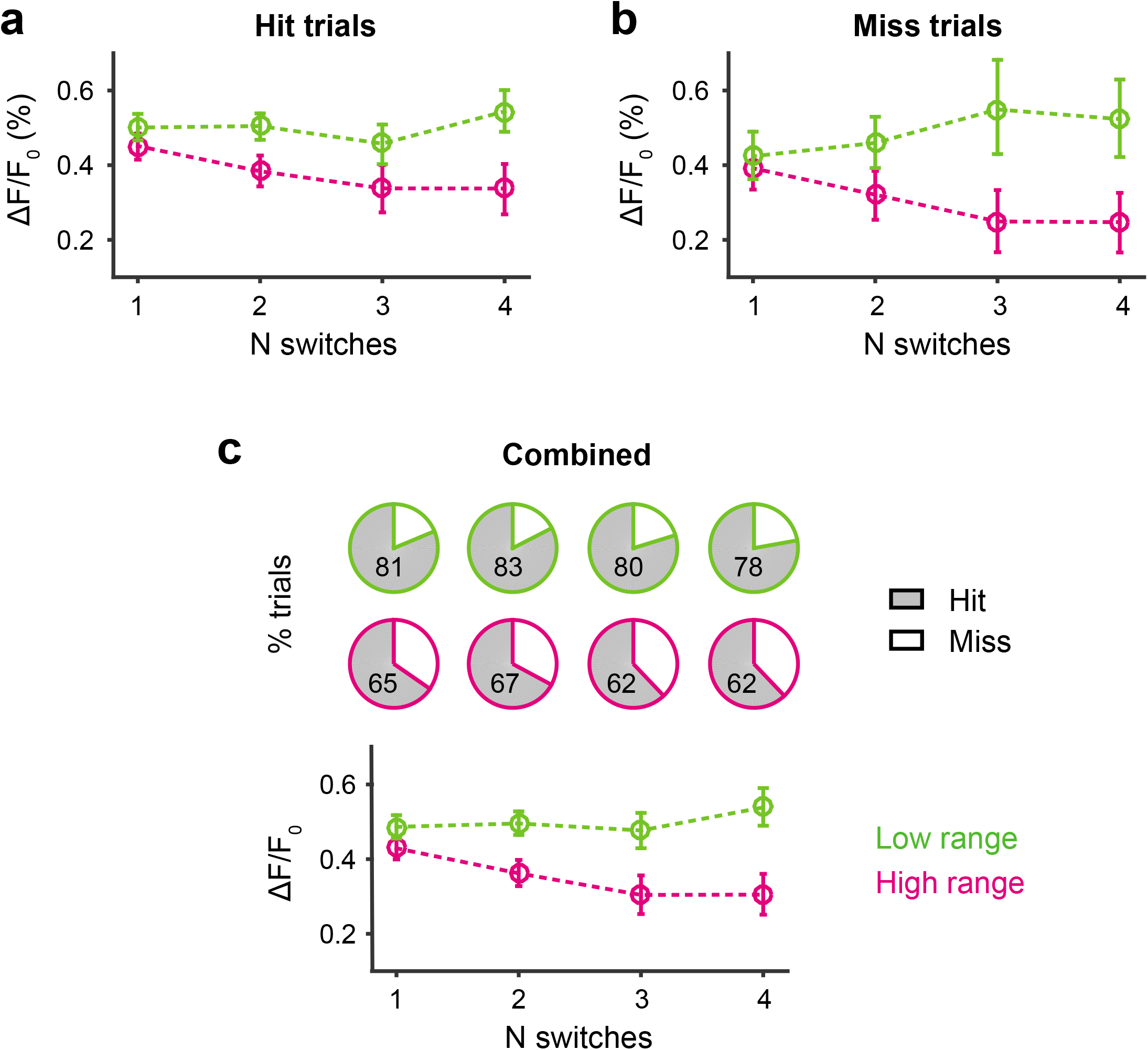
Choice-related long-term S1 dynamics. **a,** GEVI peak responses (maximum % ΔF/F_0_) for correct choices (hit trials) to 8 degree stimulation. Data is shown over multiple switches for the high- and low range condition. **b,** GEVI responses for incorrect choices (miss trials). **c,** Top: Percentage of hit and miss trials for the high- and low range condition. Bottom: Combined GEVI data (hits+misses) as a function of switches. All data shown are bootstrapped estimates of means and 95% confidence limits (based on single trials from n=6 mice, nBoot=1000 repetitions).

